# Dynamic human liver proteome atlas reveals functional insights into disease pathways

**DOI:** 10.1101/2022.01.28.478194

**Authors:** Lili Niu, Philipp E. Geyer, Rajat Gupta, Alberto Santos, Florian Meier, Sophia Doll, Nicolai J. Wewer Albrechtsen, Sabine Klein, Cristina Ortiz, Frank E. Uschner, Robert Schierwagen, Jonel Trebicka, Matthias Mann

**Affiliations:** Novo Nordisk Foundation Center for Protein Research, Faculty of Health and Medical Sciences, University of Copenhagen, DK-2200 Copenhagen, Denmark; Department of Proteomics and Signal Transduction, Max Planck Institute of Biochemistry, 82152 Martinsried, Germany; Center for Health Data Science, Faculty of Health Sciences, University of Copenhagen, 2200 Copenhagen, Denmark; Big Data Institute, Nuffield Department of Medicine, University of Oxford, OX3 7BN, UK; Center for Health Data Science, University of Copenhagen, DK-2200 Copenhagen, Denmark; Department of Clinical Biochemistry, Rigshospitalet, University of Copenhagen, DK-2100 Copenhagen, Denmark; Department of Internal Medicine I, Goethe University Clinic Frankfurt, 60323 Frankfurt, Germany; European Foundation for the Study of Chronic Failure, EFCLIF, 08021 Barcelona, Spain

**Keywords:** Clinical proteomics, tissue proteome atlas, liver disease, liver fibrosis, MS-based proteomics

## Abstract

Deeper understanding of liver pathophysiology would benefit from a comprehensive quantitative proteome resource at cell-type resolution to predict outcome and design therapy. Here, we quantify more than 150,000 sequence-unique peptides aggregated into 10,000 proteins across total liver, the major liver cell types, time-course of primary cell cultures and liver disease states. Bioinformatic analysis reveals that half of hepatocyte protein mass is comprised of enzymes and 23% of mitochondrial proteins, twice the proportion of other liver cell types. Using primary cell cultures, we capture dynamic proteome remodeling from tissue states to cell line states, providing useful information for biological or pharmaceutical research. Our extensive data serves as spectral library to characterize a human cohort of non-alcoholic steatohepatitis and cirrhosis. Dramatic proteome changes in liver tissue include signatures of stellate cell activation resembling liver cirrhosis and providing functional insights. We built a web-based dashboard application for the interactively exploration of our resource.

**Highlights:**

- Cell-type resolved liver proteome with copy numbers for 10,500 proteins
- Time-course of human liver primary cells uncovers functional proteome shifts
- A human cohort study reveals liver proteome changes in NASH and cirrhosis
- An interactive web portal integrates the results for easy exploration

## Introduction

The liver is essential for homeostasis of the human body and maintains a well-orchestrated network of parenchymal hepatocytes and multiple non-parenchymal cell types, interconnecting with the vascular and biliary system. While hepatocytes perform key metabolic functions, detoxification and protein synthesis, the non-parenchymal cells provide a microenvironment for substance exchange and promote inflammatory and immunological responses (Kmiec, 2001; Shetty *et al*, 2018). The liver is constantly exposed to gut-derived dietary antigens, viruses, bacterial products and toxic substances such as alcohol, drugs and excess lipids, all of which can induce liver damage followed by fibrosis as a repair mechanism and wound-healing response. Chronic liver injury results in persistent hepatic inflammation, which can further progress to fibrosis and eventually cirrhosis - the common end-stage of chronic liver disease (CLD). CLD – including alcoholic liver disease (ALD) and non-alcoholic liver disease (NAFLD) - is a major, global health problem affecting approximately one billion people and causing more than two million deaths annually due to complications of cirrhosis and hepatocellular carcinoma (Asrani *et al*, 2019; Loomba & Sanyal, 2013). In addition, liver disease is also important as a comorbidity, which limits or precludes effective treatment of extrahepatic diseases, e.g. in malignancy (Mokdad *et al*, 2014). Combined with its typically silent progression, there is an urgent need to implement screening programs in at-risk populations for early diagnosis (Collaborators, 2020; Ginès *et al*, 2016; Marcellin & Kutala, 2018). Existing tests are of limited performance, especially at detecting early disease stages, where they are still reversible. Thus biomarker discovery is an active research area and is ideally complemented by underlying biological mechanisms by which the marker indicates disease states. This requires knowledge of the tissue and cellular origin of the detected abnormal protein levels and the biological pathways or processes through which the protein plays a role in the disease. In this regard, we have recently investigated paired plasma and liver biopsy in a large ALD cohort, connecting circulating markers to tissue pathology, providing additional validity (Niu *et al*, 2020). However, we still lack information at cell type level which would greatly help interpret results from plasma or bulk liver tissue proteomics analysis.

Hepatocytes are the main parenchymal cells. Despite their great regenerative capacity, they may lose function, undergo fatty degeneration, release damage-associated molecular patterns (DAMPs) to maintain chronic inflammation and even transform to malignant cells during CLD. The resident hepatic macrophages (Kupffer cells) are crucial in the pathogenesis of chronic and acute liver diseases, in which they orchestrate both the resolution and progression of inflammation and tissue repair (Krenkel & Tacke, 2017; Lefere & Tacke, 2019; Wen *et al*, 2021). Hepatic stellate cells (HSCs) play a key role in the subsequent development of fibrosis (Mederacke *et al*, 2013). Upon liver injury, HSCs transdifferentiate from vitamin-A storing and quiescent HSCs to proliferative, fibrogenic, myofibroblast-like cells – a process termed HSC activation. HSCs secret growth factors, cytokines and extracellular matrix. While it is often difficult to isolate activated HSCs in patients, activation can also be induced in primary cell culture. Characterizing the proteome dynamics of this process may lead to new antifibrotic therapeutic options as most drug targets are proteins. Furthermore, such proteome shift upon primary cell culture report on the degree to which it reflects in vivo conditions (Azimifar *et al*, 2014; Heslop *et al*, 2017; Pan *et al*, 2009).

Mass spectrometry (MS)-based proteomics enables global and targeted analysis of proteins in a systematic, systems-wide and quantitative fashion (Aebersold & Mann, 2016). Although important insights into human liver proteomes has been generated by international efforts (Kampf *et al*, 2014; Sun *et al*, 2010), due to the nature of the technologies used, quantitative readout at the protein level was not sufficient to capture biological or pathological proteome dynamics.

Given the importance of liver physiology and pathology, we reasoned that dramatic advances in MS-based proteomics technologies could provide much more detailed insights into the quantitative human liver proteome. In particular, we wished to use the “proteomic ruler” approach for copy number estimation of individual proteins per cell, organelle or pathway given the fixed relation of core histones to DNA in a cell (Wisniewski *et al*, 2014), to draw quantitative cellular proteome maps. Here we performed MS-based proteomics with fractionation and label-free quantification on four major primary liver cell types derived from three individuals with normal liver histology as well as liver biopsy, and paired extrahepatic vessels (hepatic artery and portal vein) derived from six individuals undergoing liver transplantation (three healthy donors and their matched liver transplant recipients with cirrhosis). In-depth proteomic profiling and analysis revealed ‘cell-type specific’ expression patterns as well as fundamental differences in cellular proteomes which we interpret in light of anticipated functional specialization. With our extensive dataset we built a spectral library for single-shot, large-scale liver tissue proteomic analysis. Taking advantage of the phased spectrum deconvolution method (ΦSDM) (Grinfeld *et al*, 2017) and employing our MS-interfacing software called Mq.Live (Wichmann *et al*, 2019), we developed an acquisition method which enabled us to achieve deep proteome coverage in only 100 min measurement time. Applying this workflow to a human cohort of non-alcoholic steatohepatitis (NASH) and liver cirrhosis revealed dramatic proteome changes involving extracellular matrix remodeling, signaling and metabolic pathways.

We also aimed to capture proteome dynamics in cellular models of the liver. Time course measurements on primary cell culture revealed proteome shifts in a time-course fashion. When integrating results with the human study, we observed that the proteomic signatures of HSC activation largely overlap with that of liver cirrhosis, providing insights into the cellular origin of the observed proteome changes in bulk cirrhotic liver. We have developed a web-based dashboard data application that can be easily accessed by biology and clinical researchers for hypothesis generation and research verification.

## Results

### In-depth acquisition of a quantitative human liver proteome

To obtain a broad and representative overview of the liver proteome we set out to measure diverse liver cell lines, primary cells and human biopsies in great depth using the most advanced MS-based proteomics workflows. We started with four commonly used immortalized cell lines, representing three hepatic cell types (Hep G2 for hepatocytes, LX2 and TWNT4 for hepatic stellate cells and SK-Hep-1 as a proxy for liver endothelial cells). Next, we obtained primary hepatocytes (hHEPs), Kupffer cells (hKCs), liver sinusoidal endothelial cells (hLSECs) and hepatic stellate cells (hHSCs) isolated from three donors aged 57-64. Two donors had no signs of steatosis/fibrosis in the liver, and one had minimal portal inflammation (Table S1). Characteristics and purity of the isolated cells were assessed by the provider - Samsara Sciences (initial cell characterization included in Appendix). The methods they used for cell isolation have previously been shown to yield cells of high purity (DeLeve et al., 2004; Lecluyse and Alexandre, 2010). Finally, we analyzed biopsies of liver tissue, hepatic artery and portal vein from six individuals undergoing liver transplantation (three healthy donors and three cirrhotic patients undergoing liver transplantation, aged 39 to 59 years). From this total of 34 samples, we extracted peptides for MS analysis after tissue homogenization, protein denaturation, enzymatic digestion followed by eight-fold fractionation using high-pH reversed phase chromatography. We took care to completely disrupt the tissue, for adequate coverage of the extracellular matrix, and met the challenge of limited sample amount by employing a ‘loss-less’ nano-fractionator (Kulak *et al*, 2017). In this part, our aim was to build a reliable proteome atlas and we therefore analyzed each fraction by liquid chromatography-tandem mass spectrometry (LC-MS/MS) with a data-dependent acquisition (DDA) method for an in-depth proteome characterization (Fig 1a).

**Fig 1.**
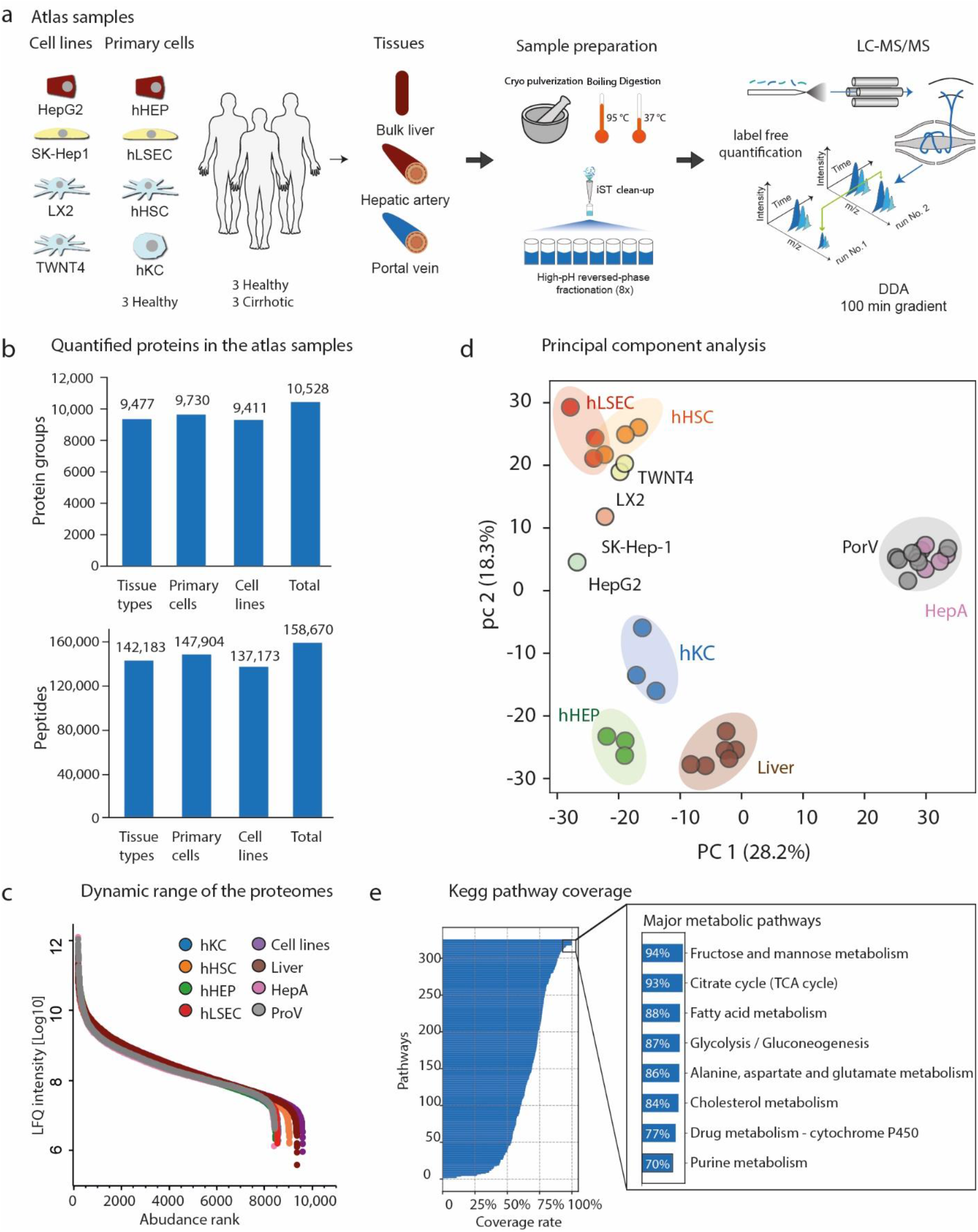
In-depth characterization of the human liver proteome. **a**. Biological material used for generating the liver proteome atlas. Samples were prepared using a bottom-up proteomics workflow followed by high-pH reversed phase fractionation, and analyzed by LC-MS/MS on an Orbitrap mass analyzer in data dependent acquisition mode. (hHSC: hepatic stellate cell, hHEP: hepatocyte, hKC: Kupffer cell, hLSEC: liver sinusoidal endothelial cell). **b**. Quantified proteome depth including the number of quantified protein groups in the upper panel and peptides in the lower panel. **c**. Dynamic range of the different proteomes based on LFQ intensity ordered by abundance rank. **d**. Principal component analysis (PCA) of all proteomes based on their proteome profiles. Where applicable replicates have the same color. **e**. KEGG pathway coverage in this dataset, with major metabolic pathways highlighted.

The resulting 272 raw data files from the 34 proteomes were analyzed using the MaxQuant software including label-free quantification with the MaxLFQ algorithm, in which we require at least two peptides with minimum seven amino acids (Cox & Mann, 2008) (Cox *et al*, 2014). Applying strict peptide and protein false discovery rates (FDR) of 1%, we identified and quantified 158,670 sequence unique peptides that were assigned to 10,528 protein groups (Fig 1b). These distinct groups sometimes resolved protein isoforms and our total identifications mapped to 9,873 protein coding genes in the human genome. Our workflow quantified 9,477 proteins in all tissue types, 9,730 in all primary cell types and 9,411 in all cell lines. Thus this most comprehensive human liver proteome data set, provides an excellent basis for systems-wide analyses (Fig 1b and Dataset EV1).

The abundance of all quantified proteins as judged by their peptide signals spans more than six orders of magnitude (Fig 1c). Analyzing the data of our proteomic effort revealed a 35% median, aggregated sequence coverage (more than two million amino acids of the liver proteome), a very high value for shotgun proteomics (Fig EV1a). The median peptide number per liver protein was 12 but ranged up to 367 for Microtubule-actin cross-linking factor 1 (MACF1, Fig EV1b).

Principal component analysis (PCA) revealed distinct proteomes between tissue types, primary liver cell types and the cell lines, with the first component capturing most of the variance (28.2%) setting apart hepatic arteries and portal veins from the rest (Fig 1d). The second component further separated hHEPs, hKCs and liver tissue from hHSCs, hLSECs and the cell lines, with 18.3% variance explained. Hepatocytes constitute about 80% of the liver cell population, therefore should be characteristic of the bulk liver proteome. Thus, hepatocytes and biopsies of the liver tissue clustered very closely, whereas hHSCs and hLSECs exhibited higher similarity to each other. LX2 and TWNT4, the two immortalized hepatic stellate cell lines grouped closely with the corresponding primary cells, whereas HepG2 was more alike to the other cell lines than primary hepatocytes, indicating proteome divergence from the in vivo states and acquisition of common cell line features.

To investigate proteomic coverage of characteristic biological pathways of the liver, we extracted the 326 pathways of the KEGG pathway database (Kanehisa *et al*, 2017; Kanehisa & Goto, 2000). For fructose and mannose metabolism, citrate cycle, fatty acid metabolism, glycolysis/gluconeogenesis and cholesterol metabolism, we obtained coverage between 84-94%, indicating that our data quantifies these processes nearly completely at the proteome level (Fig 1e). Conversely, pathways that are not expected to have biological function in the liver, say nicotine addiction and the olfactory transduction pathways, had less than 4% pathway coverage.

### Distinct proteome features between liver parenchyma and blood vessels

The liver is organized into liver lobules consisting of a portal triad, hepatocytes arranged into hepatic cords separated by adjacent sinusoids, and a central vein, and it has two major influx vessels - the portal vein and the hepatic artery. To capture the proteome differences between them, we characterized biopsies of the liver tissue, portal vein and hepatic artery. In total, 9,477 proteins were quantified across different tissues, of which 81% were common to all tissues. We exclusively quantified 824 proteins in liver biopsy but only 86 exclusive to the two blood vessels (Fig 2a), indicating both higher proteomic complexity of the liver biopsies, and the presence of blood vessels in the portal triad. When extracting the most abundant proteins in liver biopsy based on intensity of label free quantification (LFQ), we found metabolic enzymes (carbamoyl-phosphate synthase 1 (CPS1), which is heavily downregulated in NAFLD (De Chiara *et al*, 2018), alcohol dehydrogenase 1B (ADH1B) and retinaldehyde dehydrogenase 1 (ALDH1A1)), blood proteins (albumin (ALB) and hemoglobin subunits (HBA1, HBB)) and structural constituents of the cytoskeleton (vimentin (VIM), type VI collagen (COL6A3), myosin (MYH9) and keratin 18 (KRT18)) (Fig 2b). In contrast, the most abundant hepatic artery proteins were almost exclusively cytoskeleton proteins with filamin-A (FLNA), smooth muscle alpha-2 actin (ACTA2), myosin (MYH11), vimentin (VIM), type I collagen (COL1A1, COL1A2) and transgelin (TAGLN) apart from albumin and hemoglobin subunits among the top 10 (Fig 2b). Analysis of the top abundant proteins in liver biopsy and blood vessels revealed their structural and functional characteristics.

**Fig 2.**
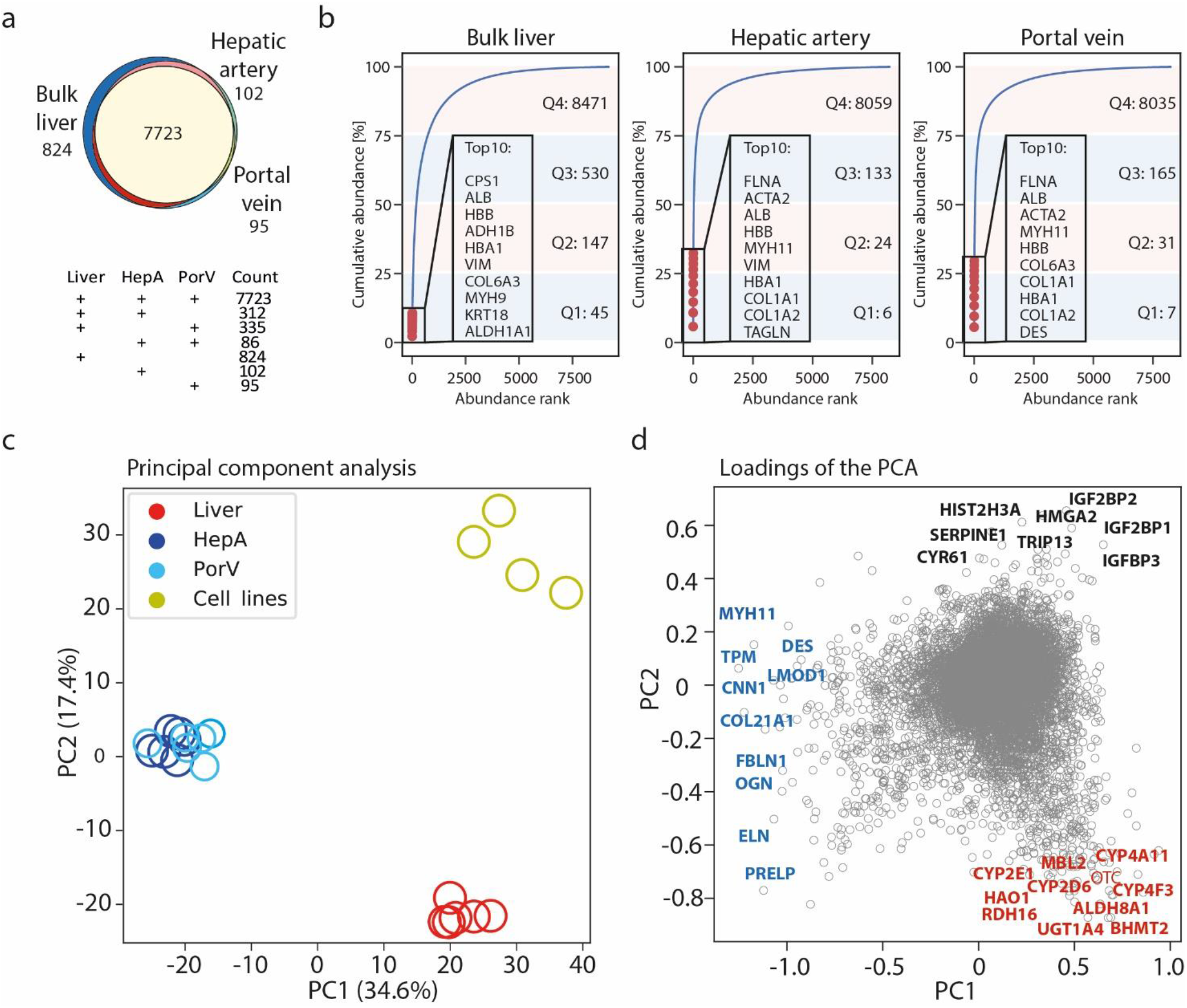
Comparative analysis of the liver tissue proteomes. **a**. Commonly and exclusively quantified proteins in bulk liver tissue, hepatic artery and portal vein. **b**. Cumulative protein abundance of liver biopsy, haptic artery and portal vein as a function of protein rank, with the total of the top ten abundant proteins and number of proteins that comprise four quartiles indicated. **c-d**. PCA of liver, hepatic artery, portal vein and cell lines with annotated proteins in the loading plot that contribute most to the variance for the three clusters.

Of note, we found that half of the total measurable proteome mass in liver is comprised of less than 200 proteins. Adding the next 523 proteins accounts for 75% of the total measurable proteome mass, whereas the remaining 25% is composed of an astonishing 8,471 proteins. The top abundant protein in liver tissue – CPS1, has a quantitative signal of more than 1 million-fold higher than the least abundant protein (ER lumen protein-retaining receptor 3, KDELR3) that we quantified. This also explains why liver is a very challenging tissue in proteomic analysis. This uneven distribution is even more pronounced in the hepatic artery and portal vein, with half of the proteome mass composed of merely 30 and 38 proteins, respectively.

Next, we investigated proteins that contribute the most to the separation of proteomes between liver biopsy and the blood vessels. In PCA, as already mentioned, biological replicates of liver biopsies and those of blood vessels clustered closely together (Fig 2c). Proteins characteristic of the blood vessels such as cytoskeleton-, and extracellular matrix proteins drive the first dimension of this separation, together with metabolic enzymes in the bulk liver tissue such as members of the cytochrome P450 superfamily: CYP2E1, CYP4A11, CYP4F3, CYP2D6, and enzymes in alcohol metabolism: ALDH8A1, RDH16, HAO1. When adding hepatic cell lines in this analysis, we found proteins involved in the regulation of cell differentiation and proliferation, such as CYR61, SERPINE1, HIST2H3A, RORC, IGF2BP1, HMGA2; regulatory proteins in cell cycle: HMGA2 and TRIP13 and proteins involved in chromatin organization: HIST2H3A, JMJD6, HMGA2 (Fig 2d). These examples illustrate how our resource can connect quantitative protein expression with functional roles of tissues and cell lines.

### Proteome differences between primary cell types reflect their functional roles

Among the non-parenchymal cells populating the liver, LSECs, hHSCs and hKCs predominate. We quantified more than 8,200 proteins in primary cells of hepatocytes and each of these cell types, and more than 7,200 across all of them (Fig 3a). As expected by their very different functions, proteome similarities were modest (average Person correlation coefficient of abundance was 0.73). The one exception were the hLSECs and hHSCs with a correlation coefficient of 0.9 (Fig 3b). The high proteome similarity between hLSECs and hHSCs in the liver has been previously observed in both proteomics and single-cell transcriptomics studies (Aizarani et al., 2019; Ölander et al., 2020). The cumulative protein abundance showed an uneven distribution similar to the bulk liver tissue.

**Fig 3.**
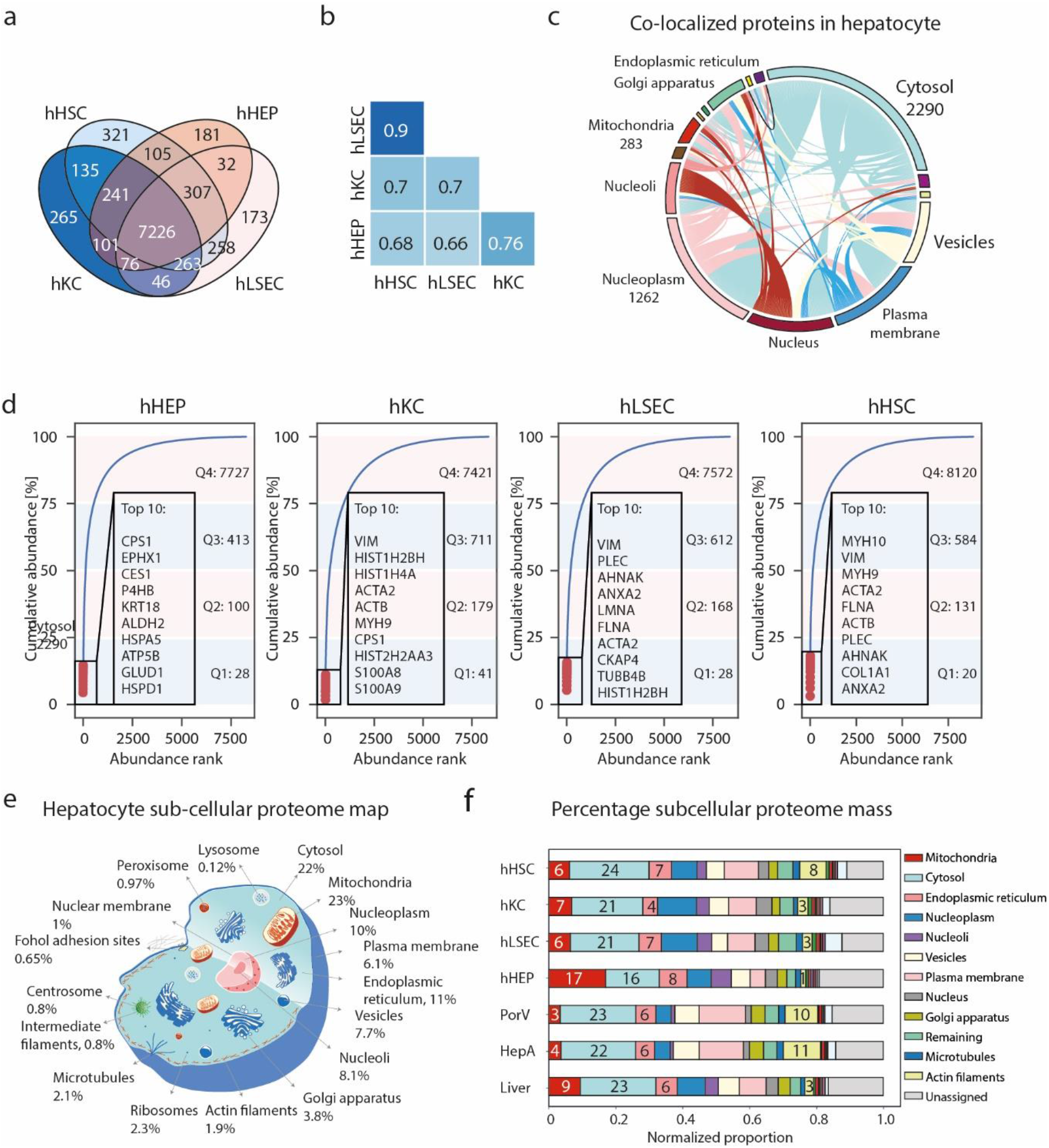
Comparative analysis of liver cell type proteomes. **a**. Commonly and exclusively quantified proteins in liver cell types (hHSC: hepatic stellate cell, hHEP: hepatocyte, hKC: Kupffer cell, hLSEC: liver sinusoidal endothelial cell). **b**. Pair-wise correlation of the proteomes of the four primary cell types, with Pearson correlation coefficients noted. **c**. Circos plot representing proteins predicated to be co-localized in subcellular compartments. **d**. Cumulative protein abundance of liver cell types as a function of protein rank, indicating the total of the top ten abundant proteins and number of proteins that comprise the four quartiles. **e**. Schematic representation of the subcellular mass composition of an average hepatocyte. **f**. The bar plot shows the contribution of each organelle to total cellular protein mass, also accounting for unassigned proteins. Percentage of cytosol, mitochondria, nucleoli, plasma membrane and actin filaments is indicated.

Interestingly, 25% of total protein mass in hepatocytes is comprised of only 28 most abundant proteins (Fig 3d). The top 10 are primarily metabolic enzymes including CPS1, liver carboxylesterase 1 (CES1), protein disulphide-isomerase (P4HB), aldehyde dehydrogenase (ALDH2), ATP synthase subunit beta (ATP5B) and glutamate dehydrogenase 1 (GLUD1). The only exceptions were heat shock proteins A5 and D1 and keratin 18, a widely used marker of hepatocyte cell death. In Kupffer cells, cytoskeleton components such as vimentin, actin isoforms and myosin are among the top abundant proteins. In the top ten most abundant proteins, they have S100A8 and S100A9, who play a prominent role in the regulation of inflammatory processes and immune response (Pruenster *et al*, 2016). Their heterodimer, termed calprotectin is also highly expressed in neutrophils and monocytes. The nature of the most abundant proteins, reflects their functional roles such as migration along the sinusoids and for the immune response when encountering infections. In both hLSEC and hHSC, again unlike in the hHEPs, the top ten abundant proteins are mostly cytoskeletal framework components, in this case presumably required for forming the endothelial fenestrae by hLSEC and for the maintenance of morphology and cellular adhesion of hHSC (Fig 3d).

The same proteins can be present in different cell types in very different amounts, depending on their functional demands. To investigate the cellular mass composition at the protein class level, we followed the annotation of proteins into 21 classes, such as enzymes, secreted protein and drug targets, of the Human Protein Atlas (HPA) (Uhlen *et al*, 2016). This revealed major differences between the cells, for example 21% of the protein mass of primary hepatocytes are classified as secreted proteins, as compared to only 12-16% in the other cell types (Fig EV2a). This is expected since a role of hepatocytes is to produce and secrete proteins into the blood. Our data show that enzymes together comprise as much as 49% of total proteome mass in hepatocytes. The number drops to 31% in hKC, and 19% in both hLSEC and hHSC. Whereas hLSEC and hHSC express higher levels of transporters, cluster of differentiation (CD) markers and ribosomal proteins compared to hHEP and hKC. In addition, hLSEC features nearly four times higher abundance of transcription factors as hHEP. Hepatocytes also have the highest abundance of FDA-approved drug target proteins (13%) and potential drug targets (24%), reflecting a focus on modulating metabolic liver functions and underlining the importance of liver disease as comorbidity (Fig EV2a). A potential use of our resource, is the knowledge of the quantitative distribution of drug targets under consideration and highlighting proteins or regulatory pathways that are present in the cell type of therapeutic interest.

To investigate the proteome composition of subcellular compartments in the liver, we mapped quantified proteins to 32 subcellular localizations according to Gene Ontology Cellular Component (GOCC) and thereby predicted a subcellular proteome map. According to this analysis, mitochondria comprise 23% of total proteome mass in hepatocytes, followed by the cytosol and the endoplasmic reticulum, with 22% and 11%, respectively (Fig 3e, Dataset EV2). As many proteins co-localize to more than one organelle (Fig 3c), we normalized the proteome proportions based on the GOCC information. This revealed that mitochondria have three times the total proteome mass in hepatocytes as in other cell types (Fig 3f, Dataset EV2). Conversely, hepatocytes have the least contribution to proteome mass from the cytoskeleton illustrated by seven times lower levels of actin filaments than in hHSCs and one thirds of those of hKCs and hLSECs (Fig 3f).

To further dissect proteome composition of mitochondria in hHEPs, we determined protein components of total mitochondrial enzymes (68%), oxidative phosphorylation (OXPHOS) complex I-IV (8.2%), ATPase (9.4%), solute carriers family 25 (5%) and mitochondria ribosomes (0.4%). These proportions were quite different in the other cell types (Fig EV2b). As an example, the summed proportion of OXPHOS complex I-IV in hepatocyte mitochondria is twice of that in hHSCs. Among the 195 solute carrier proteins quantified in our atlas data, about 15% belong to the mitochondrial carrier transporter family 25 (SLC25), together accounting for up to 6% of the mitochondrial mass. SLC25 is the largest solute transporter family in humans and central to mitochondrial function (Perland & Fredriksson, 2017; Ruprecht & Kunji, 2020). Among all 66 SLC families, SLC25 accounts for an astonishing 70% of total solute carrier proteins in hepatocytes, and 45-57% in the other cell types.

### Proteome ruler and cell-type-specific proteins

Given the pronounced differences in protein abundance across cell types at subcellular organelle- and protein class level, we were interested in estimating the absolute copy number of all the proteins. We applied the ‘proteomic ruler’ approach that uses the total histone mass to cellular DNA (Wisniewski *et al*., 2014). This concept usually assumes diploidity, whereas hepatocytes reportedly have between one and four nuclei (Thoma, 2018), therefore our copy numbers are likely to be underestimated (Fig EV3a). Our calculations resulted in two to eight billion protein molecules, corresponding to 150 to 700 pg of protein mass per cell across different cell types (Fig EV3b-c). Our rough estimation allows us to infer for instance, the stoichiometric ratios of respiratory chain subunits of associated protein complexes and ATP synthase. This shows that there are approximately 128 million protein molecules of the ATP synthase in hepatocytes, with the OXPHOS complex I to IV having 15-77 million protein copies totaling 305 million in the OXPHOS complexes I to V, up to four-fold higher than those in the other cell types (Fig 4a). tProtein subunits belonging to the same complex can be very different, illustrated by the more than 100-fold difference between the most and least abundant ATP Synthase subunits (ATP5B to ATP5S) (Fig 4b), reflecting the regulatory role of the latter as a coupling factor (Belogrudov & Hatefi, 2002; Jonckheere *et al*, 2012). Subunits alpha (ATP5A1) and beta (ATP5B) forming the catalytic core in the F1 domain of the ATP Synthase, however, have very similar copy numbers with a ratio between 0.8-0.97 (ATP5A1: ATP5B) (Fig 4b) for the four cell types, in excellent agreement with their assembly stoichiometry of 1:1 (He *et al*, 2018; Walker *et al*, 1985). The F0 membrane domain is an assembly of single-copy subunits and we found similar copy numbers between them (within two-fold difference from the mean). The only exceptions were the mitochondrial DNA-encoded ATP6 and ATP8, with less than three-fold of the mean copy number, possibly reflecting a differences in protein synthesis by cytosolic- and mitochondrial ribosomes (Table S2). Similarly, MT-CO1 and MT-CO3, two of the three mitochondrial DNA-encoded subunits in the cytochrome c oxidase complex (Complex IV), also had the lowest number of protein copies compared to the rest (ranking 12^th^ and 14^th^ among the 14 subunits). The same applies to MT-CYB in Complex III. Protein complexes consist of subunits that are produced in excess. Thus our data may point to a differential role of mitochondrial and cytosolic protein biogenesis for the OXPHOS complexes. Even at an approximate level, our estimates allow modeling the stoichiometric ratio of the overall respiratory chain machinery (Complex I-V) between different cell types based on all subunits. The data indicated that approximately 300 million protein molecules in these complexes were in hepatocytes compared to only 55 million in Kupffer cells (Fig 4a).

**Fig 4.**
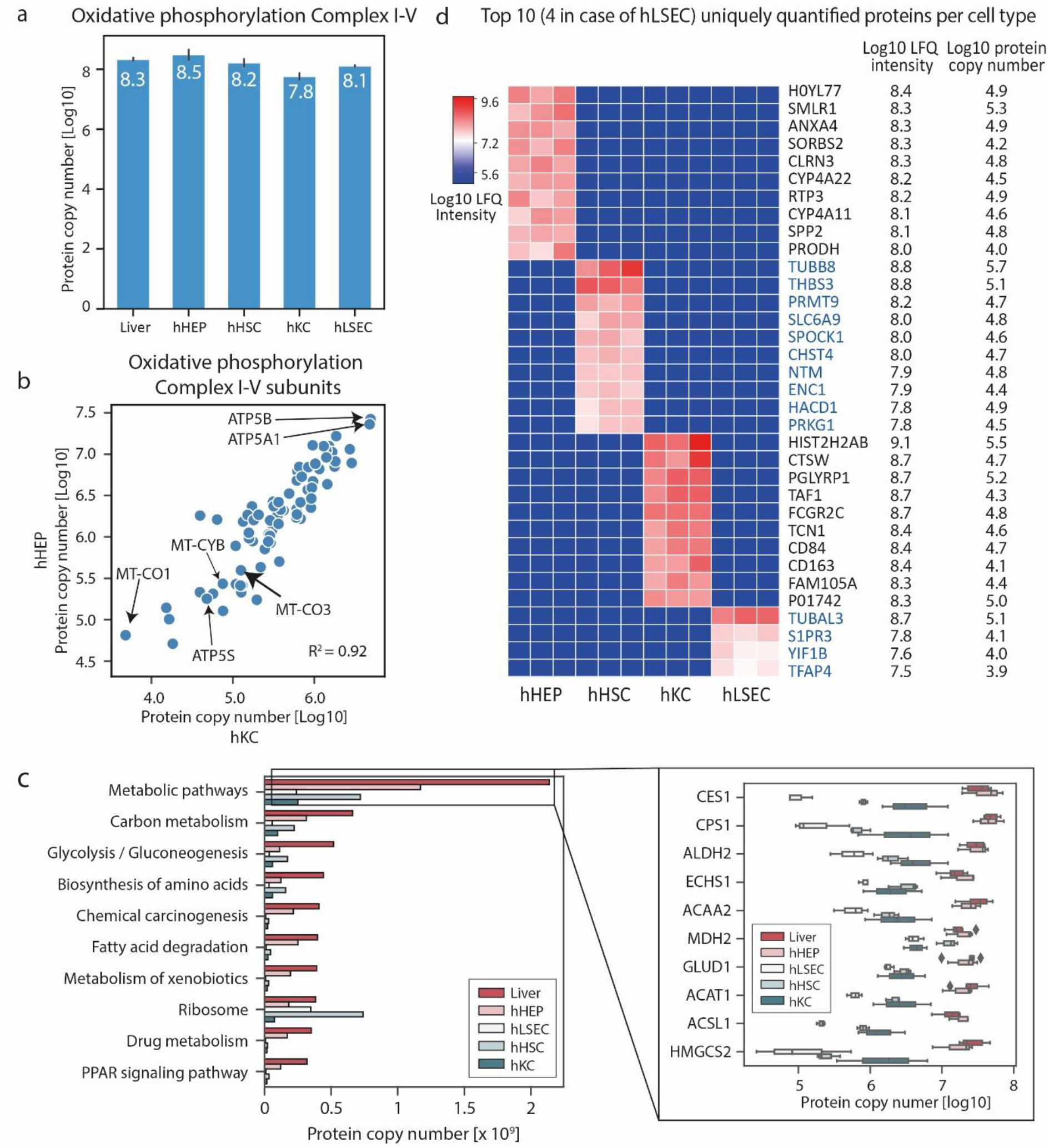
Quantitative analysis of sub-cellular proteomes and biological pathways. **a**. Total protein copy number of the oxidative phosphorylation complex I-V in bulk liver tissue and primary cell types. **b**. Protein copy number for members of the oxidative phosphorylation complex I-V in hepatocytes and Kupffer cells with Pearson correlation coefficient indicated. **c**. Protein copy number estimation of the KEGG pathways. The left panel shows the total number of proteins per cell of top ten most abundant pathways in liver biopsy in terms of total molecules of proteins associated. The right panel shows protein copy numbers of top ten most abundant proteins in liver biopsy that are associated with metabolic pathways. **d**. Top ten uniquely quantified proteins per cell type with LFQ intensities [Log10] and copy numbers [Log10].

Taking the copy number concept to another level, we extracted them for entire KEGG pathways. For instance, about 1.2 billion protein molecules in hepatocytes carry out metabolic pathway functions, whereas 240 to 720 million do so in the other liver cell types (Fig 4c). The top abundant metabolic enzymes - Liver carboxylesterase 1 (CES1) and Carbamoyl-phosphate synthase (CPS1) alone have more than 40 million protein copies each (Fig 4c).

Several genes containing non-synonymous single nucleotide polymorphisms (SNPs) have been identified to contribute to the pathogenesis of NAFLD, including PNPLA3, MBOAT7, GCKR, and HSD17B13 (Trépo & Valenti, 2020). Our copy number catalogue indicated that hHEPs express the highest of each of these proteins with the only exception of MBOAT7, which is most abundant in hHSCs (25-fold as that in hHEPs). MBOAT7 is a membrane-bound, lysophopholipid acyltransferase. Its rs641738C>T allele has been reported to be associated with fibrosis in a number of liver diseases, and it was recently shown that a loss of MBOAT7 leads to liver fibrosis, to which the mechanism is incompletely understood (Thangapandi *et al*, 2021). Our data shows that MBOAT7 has the highest expression in hHSCs, which might point to new directions in elucidating this mechanism given the prominent role of hHSCs in fibrogenesis. PNPLA3 is a lipid droplet-associated protein with hydrolase activity towards triglycerides. Individuals carrying an I148 allele on PNPLA3 has a two-fold increased risk for developing NAFLD (Romeo *et al*, 2008; Speliotes *et al*, 2010). We quantified it in only hHEPs and hHSCs with approximately 21,000 and 4,700 copies per cell, respectively. This underlines again the importance of non-parenchymal cells in development of steatosis or NASH.

TGF-beta receptor and PDGF are required for hHSC activation and inhibiting these pathways are under active investigation in terms of their anti-fibrotic potential. We further looked into their abundance levels and found that indeed, hHSCs have the highest levels of PDGF-alpha and beta (PDGFRA, PDGFRB) as well as TGF-beta receptor type-1 and type-2 (TGFBR1, TGFBR2) among all cell types, with copy numbers ranging from 150,000 to 2,000,000 (Table S3). Although, copy numbers in the other cell types are seven-fold lower on average, their higher proportion in the liver adds up to similar copy numbers. In this way, our resource provides useful information for therapeutically relevant proteins across liver cell types in relation to potential toxic effects. This builds a case why cell type-specific targeting is more effective than global approaches with less adverse effects (Klein *et al*, 2019; Klein *et al*, 2012).

Next, we investigated unique protein expression between cell types. There were only 109 proteins uniquely quantified in all the biological replicates of each cell type, and we define them as ‘cell-type specific’ (Table S4). Among the top 34 unique proteins per cell type, about 21 are among the top 5,000 abundant proteins within corresponding cell types, making it unlikely that they are uniquely detected in these cells for technological reasons (Fig 4d). Our data confirms known markers, for instance, CD163, a macrophage scavenger receptor, as a known marker of macrophages that is commonly used for Kupffer cell isolation. Three other proteins in the Kupffer cell-specific protein panel, namely peptidoglycan recognition protein 1 (PGLYRP1), cathepsin W (CTSW) and CD84 are involved in the immune response. Apart from the liver, these proteins also have ‘enriched expression’ according to RNA-seq data in the HPA in bone marrow and lymph nodes, in agreement with the similarity of Kupffer cells as resident macrophages with the infiltrating macrophages from the bone marrow before they migrate to the liver. Among the top ten hHEP-specific proteins, six have ‘enriched expression’ in the liver, namely small leucine rich protein 1 (SMLR1), clarin 3 (CLRN3), members of the cytochrome P450 family (CYP4A22, CYP4A11), receptor transporter protein 3 (RTP3) and secreted phosphoprotein 2 (SPP2) with protein copy numbers up to 180,000 (Table S4). RTP3 and SPP2 are ‘exclusively expressed’ (meaning exclusively detected) in the liver according to the HPA to which our data assigns quantitative and cell type resolved information, with more than 70,000 and 50,000 protein copies per cell, respectively (Fig 4d). A recent study has identified CLRN3, as a cell-surface protein for hepatocyte-like cells derived from induced pluripotent stem cells (Mallanna *et al*, 2016). Interestingly, the top hHEP-specific protein is an uncharacterized protein (H0YL77). Apart from hepatocytes, it was also quantified in the bulk liver samples and the HepG2 cell line, demonstrating its specificity to hepatocytes. When calculating the Pearson correlation coefficients of its abundance profile with other proteins, the highest correlation was to mitochondrial fission factor (MFF) (r=0.99) (Table S5).

Among the top ten hHSC-specific proteins, the HPA assigns three as primarily expressed in the cerebellum or cerebral cortex, namely testican-1 (SPOCK1), neurotrimin (NTM) and ectodermal-neural cortex 1 (ENC1). It has been hypothesized that HSCs derived from the neural crest due to their similar gene expression pattern to that of neural cell types (Sato *et al*, 2003), which our data supports at the quantitative level for these uniquely expressed proteins. Note that the neural crest origin of HSC has been challenged, and another hypothesis suggests multipotent mesenchymal progenitor cells as the origin, particularly for these cells also give rise to neural cells and mesenchymal lineages (Friedman, 2008; Hellerbrand, 2013). In line with the fundamental role of HSCs in producing extracellular matrix components and the initiation, progression and regression of liver fibrosis (Mederacke *et al*., 2013), we found carbohydrate sulfotransferase 4 (CHST4) among their top ten specific proteins. This protein plays an important role in lymphocyte homing at sites of inflammation. It has been linked to liver disease and was shown to predict the prognosis of hepatocellular carcinoma (Hu *et al*, 2021; Zhang *et al*, 2020). Only four proteins were hLSEC-specific by our criteria, likely due to their proteome similarity with hHSC. The four proteins are tubulin alpha chain-like 3 (TUBAL3), transcription factor AP4 (TFAP4), Yip1 interacting factor homolog B (YIF1B) and sphingosine-1-phosphate receptor 3 (S1PR3) (Fig 4d). Together, our analysis identifies known as well as so far undescribed cell-type specific proteins in the liver and provides their abundance levels.

### Functional specialization of human liver cell types

We further investigated the functional characteristics of the different liver cell types in an unbiased manner using ANOVA, which showed that nearly half of the liver proteome was significantly different in at least one of them (4,173 proteins, Methods, Dataset EV3). After hierarchical clustering based on Euclidean distance, four large clusters of cell type characteristic proteins appeared (1,000, 900, 1,168 and 1,331 proteins enriched in hHEPs, hKCs, hLSECs and hHSCs, respectively, Fig 5a). These panels are very distinct between hHEPs and hKCs but overlap between hLSECs and hHSCs. Reflecting its active role in energy metabolism and maintaining homeostasis, the corresponding GO terms are highly and significantly enriched in hHEPs, along with complement activation, blood coagulation, and regulation of blood lipoprotein levels (Fig 5b). Higher protein levels of the oxidative phosphorylation machinery in hepatocytes from the above targeted analysis was also reflected in this unbiased analysis (Fig EV4).

**Fig 5.**
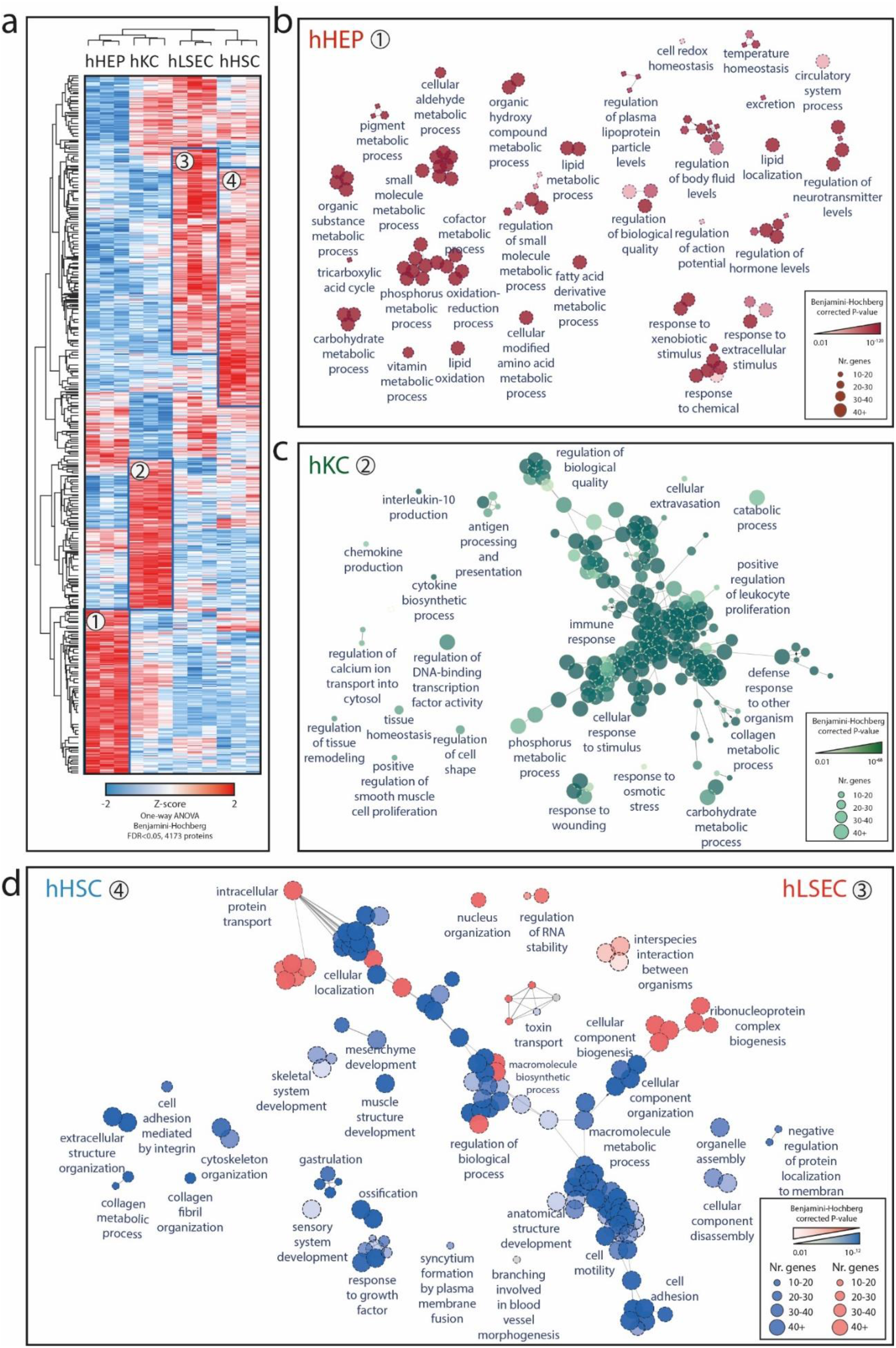
Liver cell type functional specialization map. **a**. Unsupervised hierarchical clustering of significant proteins after One-way ANOVA analysis of LFQ protein abundance across cell types, with columns showing three biological replicates of four cell types and rows significant proteins. Frames and numbers indicate four clusters of proteins highly enriched in each cell type. **b**. GOBP map specific for hHEP (proteins in cluster 1). Each circle represents an individual biological process term significantly enriched, with the number of proteins and FDR-corrected p-value indicated by size and degree of transparency. The leading term given by the most proteins associated within a group is indicated. **c**. GOBP map specific for hKC (proteins in cluster 2). **d**. Comparative analysis of GOBP enrichment for hHSC and hLSEC from Cluster 3 and Cluster 4 from Fig 5a, with blue terms specific for hHSC and pink specific for hLSEC and gray terms shared between them.

As expected, GO terms associated to immune responses were highly enriched in hKCs, including antigen presentation and processing, cytokine and chemokine production, as well as cell motility, possibly required for its movement along the liver sinusoids (MacPhee *et al*, 1992). Surprisingly, several metabolic pathways were also significantly enriched, including phosphorus metabolic process, monosaccharide metabolic process and collagen metabolic process. This is demonstrated by an higher abundance of dehydrogenases e.g. GPD1L (glycerol-3-phosphate dehydrogenase 1-like protein), GAPDH (glyceraldehyde-3-phosphate dehydrogenase), ME1 (NADP-dependent malic enzyme); carbohydrate kinases (PGK1 (phosphoglycerate kinase 1), PFKL (ATP-dependent 6-phosphofructokinase) as well as ALDOA, a fructose-bisphosphate aldolase known to be predominantly expressed in skeletal muscle.

Proteases, such as cathepsins (CTSG, CTSB, CTSD, CTSH, CTSL, CTSS, CTSW and CTSZ) which are proteolytic enzymes that contribute to pathogen killing in lysosomes (Pires *et al*, 2016), proteins of the core machinery of ubiqutination and proteasomal degradation and matrix metalloproteinases (MMP8, MMP9, MMP25) that have crucial regulatory functions have their highest abundance in Kupffer cells. Further reflecting the roles in engulfing pathogens into lysosomal compartments to undergo degradation pathways, vacuolar protein sorting-associated proteins involved in cargo transport, such as VPS11 and VPS26A were most highly expressed compared to the other cell types (Fig EV5).

As most hHSCs- and hLSECs-enriched proteins overlapped, we did a comparative functional enrichment analysis on proteins from cluster three and four (Fig 5a). Enriched terms were termed dominant in a cell type if the percentage of associated proteins from the cell type is larger than the other. Among the 26 terms specific for hLSECs, a considerable proportion relates to ribonucleoprotein complex biogenesis, gene expression and RNA localization. Hence, levels of 40S- and 60S- ribosomal subunits as well as 39S and 28S mitochondrial ribosomal proteins, members of the nuclear pore complex, RNA helicase, and translation initiation among the others where comparatively higher, reflecting higher biosynthetic activity due to the important role of LSEC in liver homeostasis, especially since LSEC is the first barrier of nutrients and ‘pathobionts’ entering the human body through the gut.

LSEC constitutes the sinusoidal fenestrae, and plays an essential role in the exchange of solutes, metabolites, fluid and particles between the hepatocytes and sinusoidal blood (Knolle & Wohlleber, 2016). Accordingly, proteins involved in intracellular transport such many SLC family members were highly abundant compared to other cell types (Fig EV6). Membrane proteins with virus receptor activity involved in interspecies interaction were also highly enriched such as CD46, ICAM1, PVR (poliovirus receptor), EPHA2 (ephrin type-A receptor 2, which acts as a HCV receptor in hepatocytes and facilities its entry), as well as ANPEP (also known as CD13), DPP4 (also known as CD26) which both have human coronavirus receptor activity (Peck *et al*, 2017; Sungnak *et al*, 2020; Tang *et al*, 2020), presumably rendering LSEC more sensitive or susceptible to viral infection or translocation through receptor-mediated endocytosis or fusion. Among the HSC-specific enriched terms, many relate to anatomical structure development and extracellular structure organization (Fig 5d). Highly abundant cytoskeleton component proteins such as actin and tubulin, motor protein myosin, membrane-cytoskeletal protein vinculin, transmembrane receptor integrins and extracellular matrix proteins (laminin and collagens) all belong to this category. Alpha-smooth muscle actin (α-SMA) is a marker for HSC activation but its expression is not unique to HSC, with up to ten-fold lower copy numbers in the other cell types in our resource.

### Primary cell culture reveals hepatic stellate cell activation-related proteome signatures

Functional studies related to the liver and its cell types are mostly performed in primary or immortalized cell lines due to convenience and reproducibility. We previously compared immortalized murine hepatocyte cell lines to their cognate primary cells, which revealed rearrangement of characteristic metabolic processes, decrease in mitochondria function, while insulin signaling remained intact (Pan *et al*., 2009). In mouse tissue and isolated hepatocytes, expression of complement components gradually decreased as one aspect of extensive overall proteome remodeling during cell culture (Azimifar *et al*., 2014). In another proteome study, a decrease in expression of cytochrome P450 family was observed in human hepatocyte cell culture (Heslop *et al*., 2017).

Our liver proteomics workflow could shed light on which cellular functions are retained in human liver cells during adaption from an in-vivo like state to a cell line state. To test this hypothesis, we performed primary cell culture of hHEPs, hHSCs, and hLSECs and investigated their proteomic changes at day-1, day-3, and day-7. During the cell culture experiment, we chose optimal culturing conditions for each primary cell type according to the provider’s instructions and systematically controlled the viability and confluency to avoid introducing proteome changes due to sub-optimal growth conditions (Methods). We acquired the mass spectra in data-independent acquisition (DIA) mode with FAIMS, and in-silico library (directDIA) was used for data processing (Bruderer *et al*, 2015) (Methods). In biological triplicates and single-run proteomic measurements, 7,501 proteins were quantified in total (Fig 6a, Dataset EV4). Hepatocytes had the largest proteome alterations comprising 41% of the total in hHEP (2,531 proteins, ANOVA, FDR<0.05) followed by hHSC (13%) and hLSEC (3%) (Dataset EV4). Despite this proteome drift, the same cell types still cluster in PCA, with the separation of hHEPs from hHSCs and hLSECs in the first component already explaining 57% of the variance (Fig 6b). This recapitulates the data from uncultured primary cells as described above, in which the proteomic profiles of hHSC and hLSEC are most alike. The PCA clearly shows the step-wise proteome shift of these primary cells from day-1 to day-7, which was further reflected by decreased Pearson correlation coefficients (0.91 from day-1 to day-3 and 0.8 from day-1 to day-7 in case of hHEP) (Fig 6c).

**Fig 6.**
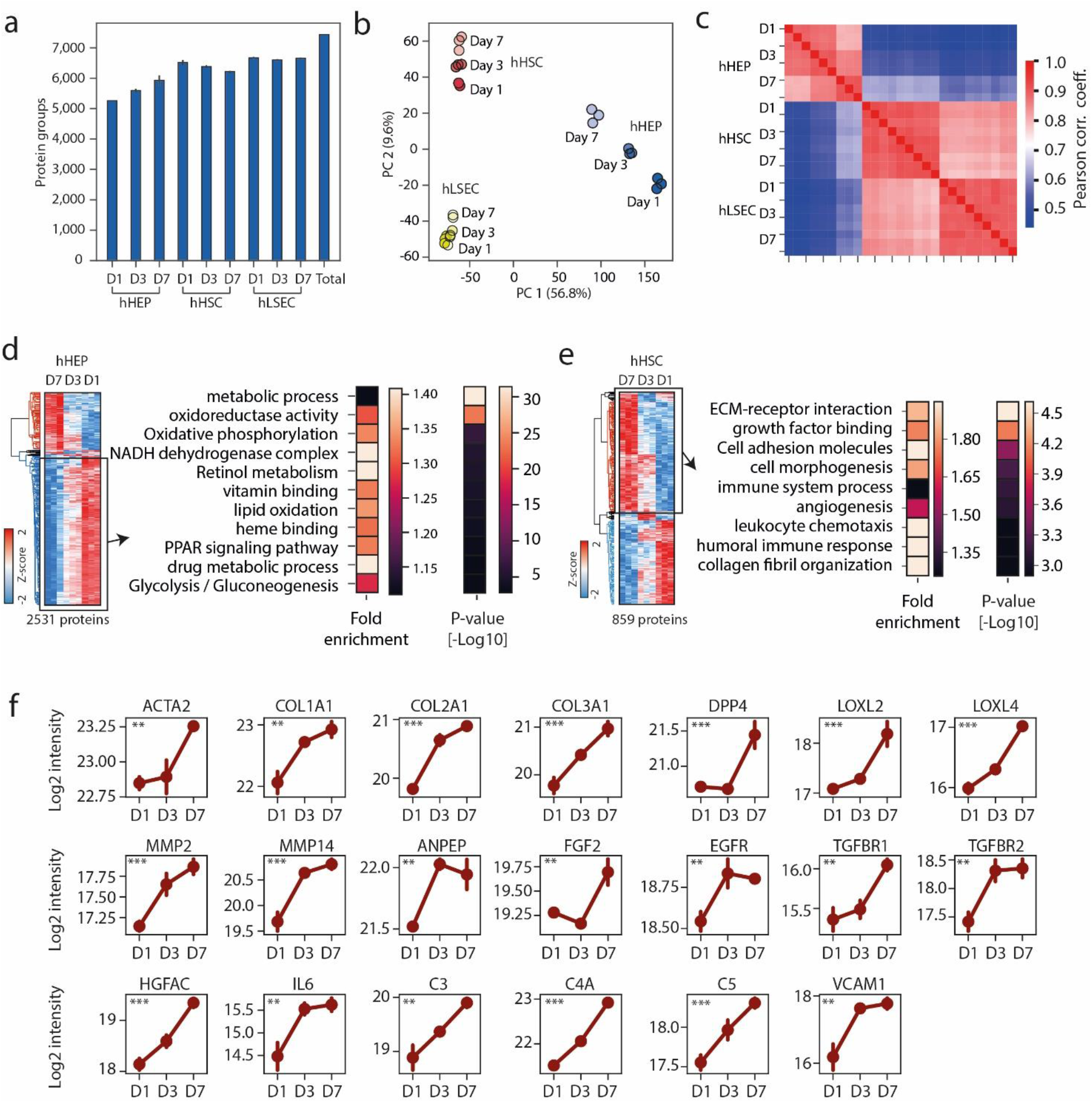
Proteomics analysis of human liver primary cells upon cell culture. **a**. Total quantified proteins at day one, three and seven upon primary cell culture in hepatocytes (hHEP), hepatic stellate cells (hHSC) and liver sinusoidal endothelial cells (LSEC). **b**. Principal component analysis showing proteome drift of liver primary cells upon cell culture. **c**. Pair-wise correlation of proteomes during cell culture. **d-e**. Significantly changing proteins during primary hHEP (d) and hHSC (e) cell culture, indicating significantly enriched GO terms in the down-regulated (hHEP) and up-regulated (hHSC) cluster. **f**. Protein expression patterns over the course of primary hHSC culture with mean and standard deviation presented. These proteins belong to the ECM, ECM-related enzymes, growth factors, and the immune system. Significance was defined by ANOVA followed by Benjamini–Hochberg correction for multiple hypothesis testing with a significance level of *P < 0.05, **P < 0.01, and ***P < 0.001.

To understand the proteome shifts at the level of biological processes, we performed functional enrichment analysis of the differentially abundant proteins. For hHEP this unveiled an upregulation of proteins involved in regulation of cell shape, adhesion and migration, as exemplified by actin and actin-binding proteins (myosin, tropomyosin, gelsolin, drebrin) as well as extracellular matrix proteins (VWF, tenascin-X, HSPG2, collagen type III, IV and VI), presumably reflecting loss of the supporting in vivo structural framework as well as mechanical adaption to the in vitro environment. Conversely, metabolic or energy homeostatic processes were downregulated, accounting for half of the altered proteome, including more than 70 members of the oxidative phosphorylation machinery components. Likewise, we observe a downregulation of GO terms related to binding vitamins and metal, along with PPAR (peroxisome proliferator-activated receptor) signaling.

PPARs are key metabolic regulators, whose agonists are therapeutic targets for NAFLD/NASH currently under evaluation in phase I to III clinical trials, including the drug Pioglitazone (Boeckmans *et al*, 2019; Della Pepa *et al*, 2021; Wu *et al*, 2020). The time-dependent decrease in PPAR emphasizes the importance to take proteome drift into account when evaluating PPAR agonist efficacy in hepatocyte models. This also applies for glycolysis/gluconeogenesis, lipid oxidation and drug metabolic processes, which all decrease upon cell culture (Fig 6d).

Interestingly, hHSCs underwent gradual loss of vitamin A-containing lipid droplets during primary cell culture – a typical feature of HSC activation. We confirmed this with our proteomics data, illustrated by an upregulation of the HSC activation marker α*-*SMA by 30% at day-7 compared to day-1 (Fig 6f). HSC activation is a key event in fibrogenesis, during which quiescent HSCs differentiate into myofibroblast-like cells and secret excessive extracellular matrix. This process is crucial for understanding the pathogenesis and development of liver fibrosis. Functional analysis of the more than 800 significantly changing proteins furthermore revealed increased expression in collagen microfibril organization, extracellular matrix organization and angiogenesis (Fig 6e). Specifically collagen type I, II, III, ECM1, as well as extracellular matrix modifiers Lysyl oxidase (LOXL2, LOXL4) had an about two-fold increase at day 7 compared with day 1. Among these, LOXL2 has emerged as an attractive therapeutic target for inhibiting liver fibrosis (Barry-Hamilton *et al*, 2010; Ikenaga *et al*, 2017). The collagenase MMP2, MMP14 and their inhibitors TIMP2 which also have anti-fibrotic therapeutic potential were as well upregulated by 65%-120%, likely reflecting a higher turnover rate of extracellular matrix (Chuang *et al*, 2019; Craig *et al*, 2015).

Several receptor tyrosine kinases and related proteins, known to be implicated in hepatic fibrosis or liver regeneration - including Epidermal growth factor receptor (EGFR), Fibroblast growth factor 2 (FGF2) and Hepatocyte growth factor activator (HGFAC) – were also upregulated (Fig 6f). Liver disease such as cirrhosis can result in changes in the growth hormone-insulin-like growth factor axis (Bonefeld & Møller, 2011; Donaghy *et al*, 2002). In line with this, we observed about 50% higher levels of insulin growth factor binding proteins (IGFBP3, IGFBP4, and IGFBP7) by day-7 upon HSC activation. Transforming growth factor (TGF)-β is a major profibrogenic cytokine and targeting TGF-β signaling has been explored to inhibit liver disease (Breitkopf *et al*, 2005; Rao & Mishra, 2019). Accordingly, TGF-β receptors TGFBR1 and TGFBR2 levels were also more than 50% higher on day-7 during HSC activation.

In previous studies we had found that the peptidases ANPEP and DPP-4, a well-known drug target for T2D, were associated with NAFLD in a human cohort and mouse models (Niu *et al*, 2019), and we found them to be significantly and substantially upregulated upon HSC activation (Fig 6f). Furthermore, we had employed unbiased machine learning algorithms to select a panel of 14 circulating markers for predicting fibrosis in alcohol-related liver disease (Niu *et al*., 2020). Levels of four of these proteins – all among the top predictors of fibrosis in ALD - increased 1.5 to 3-fold, namely VCAM1, IGFBP7, IGFALS, and LGALS3BP. Thus, our dynamic proteomic profile of HSC activation provides functional insights into liver fibrosis and can allocate cellular origin of circulating markers.

Interestingly, ‘immune system process’ and ‘leukocyte chemotaxis’ were among the most highly enriched GO terms in upregulated proteins during cell culture of primary hHSCs (Fig 6e). The depth of our experiments allowed the identification of IL-6, a protein of extremely low abundance in the plasma and generally considered to be secreted by macrophages such as the liver resident Kupffer cells and modulate hepatic inflammation (Han *et al*, 2020; Schmidt-Arras & Rose-John, 2016). Its levels doubled already by day-3 of HSC culture. Altogether, more than 70 immune response proteins were upregulated, including the complement components (C3, C4A, C5, C8B) and proteins involved in regulation of leukocyte chemotaxis (VCAM1) (Fig 6f). Thus, apart from ECM remodeling, HSC activation may perpetuate hepatic inflammation by secreting pro-inflammatory factors to recruit leukocytes to the liver. These findings revealed a potential unappreciated role of HSC activation in the development of fibrogenesis.

### Dramatic proteome landscape shift in cirrhotic liver

Having generated an in-depth human liver proteome, we used it to build a ‘spectral library’, to facilitate single-run DIA analysis of liver biopsies in clinical cohorts. We applied this rapid and sensitive pipeline to investigate and compare various pathological conditions of the liver. Investigated liver samples were from cirrhosis patients requiring liver transplantation as the maximal end-stage of CLD (N=10), patients with morbid obesity and non-alcoholic steatohepatitis requiring bariatric surgery (NASH, N=20), and 15 healthy controls, of which 10 were obese but liver-healthy. Participant characteristics can be found in Table S6. In total, we quantified 6,475 protein groups (Dataset EV5). After filtering for at least 70% valid values in experimental groups, we obtained a data-matrix with an average 5,382 proteins per sample and an overall data completeness of 91.8%) (Methods). ANOVA analysis resulted in 2,772 proteins differentially expressed between cirrhosis, NASH and healthy controls (Fig 7a, Dataset EV5). Of these, two thirds had increased abundance in liver cirrhosis and they are involved in extracellular matrix remodeling, signal transduction, cell morphogenesis and migration, immune response and angiogenesis (Fig 7b, Dataset EV5). These results provide the molecular basis for the clinical observations of liver cirrhosis, characterized by the replacement of normal liver tissue by scar tissue, the formation of new vessels leading to abnormal angioarchitecture in the cirrhotic liver, and the compromised immune system and dysregulated immune cell activation (Schierwagen *et al*, 2020; Trebicka *et al*, 2019). Both TGF-β signaling and platelet derived growth factor signaling play an important role in mediating hepatic stellate cell activation and development of fibrosis. We found them to be upregulated in cirrhosis represented by the upregulation of PDGFRA, PGDFRB, TGFB1 (TGF-β1), TGFB1I1 (TGF-β1 induced transcript 1 protein), TGFBI (TGF-β induced protein ig-h3), LTBP1, LTBP4, and ENG (transmembrane accessory receptor for TGF-beta signaling) (Fig 7c).

**Fig 7.**
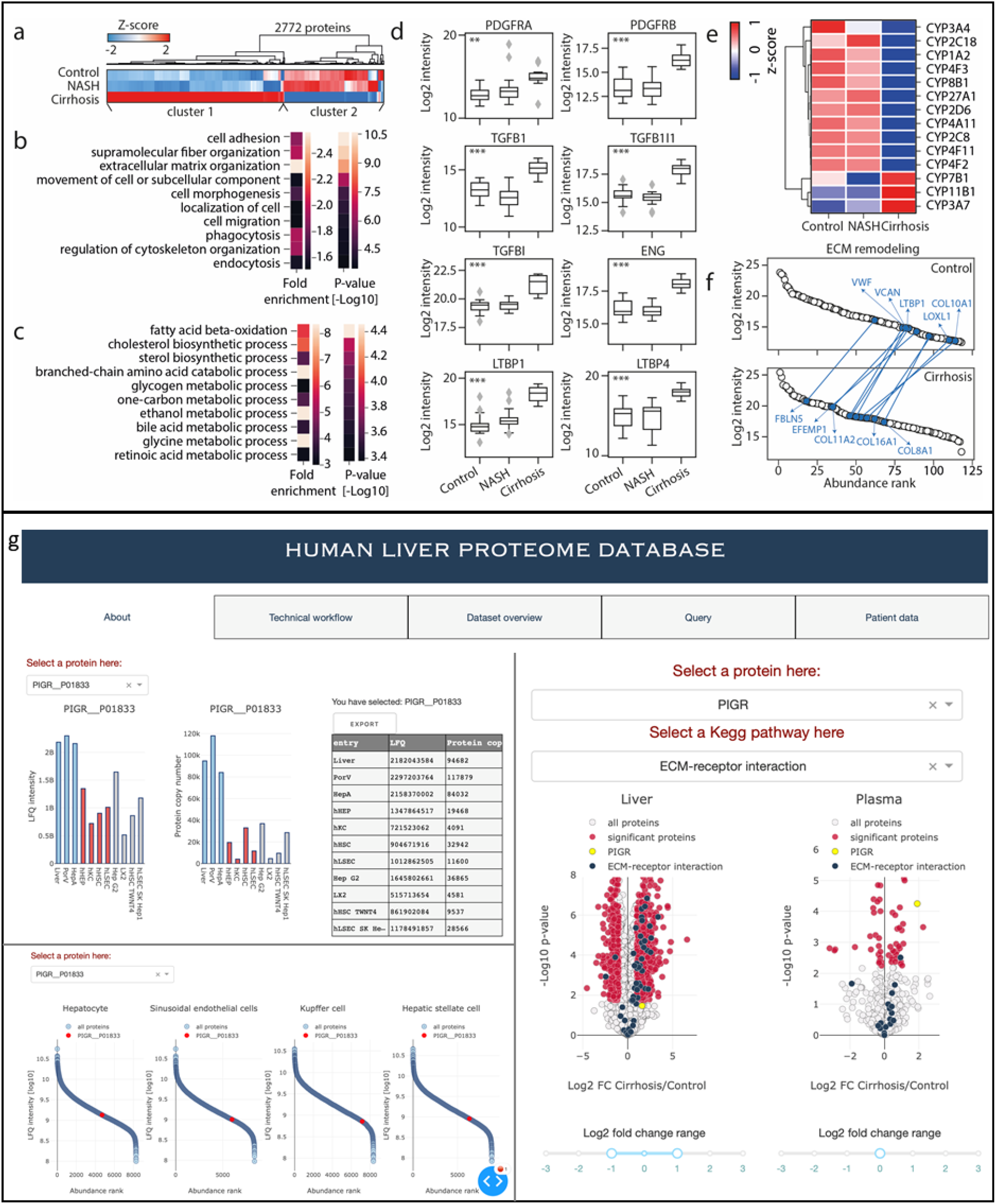
Liver proteome remodeling under pathological conditions. **a**. Hierarchical clustering of proteins significantly differentially abundant between NASH, cirrhosis and control groups (ANOVA, FDR<0.05). Two major clusters of proteins were identified with Cluster 1 mainly upregulated in cirrhosis compared to NASH and controls and Cluster 2 downregulated. **b**. Ten representative highly enriched GOBP terms in proteins in Cluster 1 of Panel (a). **c**. Box-and-whisker plot showing the distribution of log2-intensity values of statistical significantly regulated proteins across three groups (ANOVA, FDR<0.05). Number of replicates is n=15, 20 and 10 for the control, NASH and cirrhosis group, respectively. The black line in the middle of the box is the median, the top and the bottom of the box represent the upper and lower quartile values of the data and the whiskers represent the upper and lower limits for considering outliers (Q3+1.5*IQR, Q1-1.5*IQR) where IQR is the interquartile range (Q3–Q1). Significance was defined by ANOVA followed by Benjamini–Hochberg correction for multiple hypothesis testing with a significance level of *P < 0.05, **P < 0.01, and ***P < 0.001. **d**. Ten representative highly enriched GOBP terms in proteins in Cluster 2 of Panel (a). e. Heatmap of CYP 450 family members that are statistically significantly abundant between three groups. Mean protein abundance followed by Z-score normalization across groups is presented. Number of replicates is same as Panel (c). **f**. Extracellular matrix (ECM) remodeling in liver cirrhosis. Upper and lower panel shows the abundance rank of proteins involved in ECM organization in the control (upper) and cirrhosis (lower) groups, highlighting top shifted ECM components in the cirrhosis group as compared with the control group. **g**. A web-based dashboard app incorporating the data generated in this study.

In contrast, down-regulated proteins in liver cirrhosis are related to fatty acid metabolism, ethanol/drug metabolism and retinol/retinoid metabolic processes reflecting metabolic impairment in patients with liver cirrhosis (Fig 7d). Decreased level of retinol/retinoid metabolic process likely reflect the loss of lipid-storing phenotype, in particular the loss of retinyl ester-containing lipid droplets in HSC – a key feature of HSC activation. Interestingly, while majority of the CYP450 family enzymes are downregulated as expected, there are three exceptions – CYP3A7, CYP7B1 and CYP11B1 (Fig 7e). CYP3A7 is found predominantly in human fetal livers and is the major hepatic enzyme of the CYP450 family enzymes in the fetus (Li & Lampe, 2019). We detect an 18-fold increase of CYP3A7 in cirrhotic liver compared to controls. This unexpected finding was consistent with a transcriptomics study in which expression of CYP3A7 was found to be slightly higher in HBV cirrhosis compared to normal livers while other CYP3A family members generally decrease (Chen *et al*, 2014). CYP7B1 is a crucial enzyme in the alternative bile acid synthetic pathway (Jia *et al*, 2021), and it has not been related to fibrosis. Similarly, little is known about the expression of CYP11B1 in liver cirrhosis, and we detect it to be upregulated by 77%. Thus, this study would add to our understanding of the CYP450 family members in liver diseases and provide new research directions for the pathogenesis and progression of liver diseases.

Hepatic fibrosis is characterized by excess accumulation and dynamic remodeling of ECM. Proteomics has the ability to comprehensively characterize the ECM molecular composition and its quantitative changes in liver fibrosis, which is essential for gaining insights into the mechanisms of liver disease. The altered ECM landscape in liver cirrhosis include the upregulation of collagens (type I, III, IV, V, VI, VIII, X, XI, XII, XIV, XV, XVI, XVIII, XXI), proteoglycans such as versican, decorin, lumican, and glycoproteins including fibulins, fibronectin and laminins (Dataset EV5). Type X and XI collagens (COL10A1 and COL11A2) are the highest up-regulated collagens, even though type I collagen (COL1A1, COL1A2) is the most abundant protein in the ECM, suggesting that not only the overall abundance but also the quantitative composition of the ECM constituents is altered in cirrhotic liver (Ortiz *et al*, 2021; Praktiknjo *et al*, 2018). To investigate this in a quantitative manner, we extracted all significantly elevated ECM associated proteins in cirrhotic liver and plotted their abundance rank in cirrhotic and healthy liver respectively which revealed 20 proteins whose abundance rank shifted by at least 20 (Fig 7f). COL10A1 had the most dramatic shift – from 114 in healthy control to 53 in cirrhotic liver, followed by EFEMP1, LOXL1, COL11A2 and FBLN5. In healthy liver, the ECM is constantly undergoing remodeling processes, by which components are being modified and degraded, tightly controlled to ensure homeostasis. Among the up-regulated proteins associated with ECM organization were the matrix metalloproteinases MMP2, MMP14, MMP23A, ADAMTS5 and their tissue inhibitors (TIMP1, 3), as well as lysyl oxidase (LOXL1) which catalyzes collagen cross-link formation, indicating increased collagen crosslinking. Interestingly, many of the overexpressed proteins such as collagens, Lysyl Oxidase, tissue metalloproteinase inhibitors were also upregulated upon hepatic stellate cell activation, indicating their potential cellular source.

Unlike the dramatic proteome shift in cirrhotic liver, NASH featured only marginal changes in liver proteome compared to normal livers characterized by 325 proteins significantly differentially abundant (Tukey post hoc test on ANOVA significant hits with FDR<0.05) (Dataset EV5). NASH resulted in less proteome changes in the liver compared to cirrhosis, consistent with a transcriptomics study, in which only dozens of significantly differentially expressed proteins (DEPs) were detected in NASH and more than 1,000 DEPs in cirrhosis (Govaere et al., 2020).

To provide an open interface for easily accessing the data resource generated in this study, we built a web-based Dashboard app that enables interactive data exploration and exportation (Methods, Fig 7g). The database provides intuitive ways of data inference, such as (i) proteome-wide inference of protein abundance across liver tissue and cell types with both MS intensity and protein copy number, (ii) inference of protein abundance rank in a cellular proteome across four primary liver cell types, and (iii) inference of proteome changes in liver and plasma upon pathological conditions such as liver cirrhosis at protein and pathway level. In addition, the database provides information of observed peptides for each identified protein, which can be useful for building targeted assay such as parallel reaction monitoring.

## Discussion

The liver is a vital organ responsible for hundreds of functions in the body. Multiple risk factors predispose to liver diseases, which impose a huge burden on global health systems, affecting an estimated one billion people (Loomba & Sanyal, 2013). A better understanding of basic liver biology and the underlying mechanism of pathology can aid prevention, diagnosis and treatment. In the era of “big data” when omics data are increasingly being generated in human cohorts, a fast and sensitive proteomics workflow for liver tissue is required.

MS-based proteomics is a constantly developing field. Through strategies such as multi-enzyme digestion, extensive fractionation, the use of various fragmentation techniques, and matching to other tissue proteomes, current technology has been able to identify >10,000 proteins in bulk human liver tissue at the expense of analysis time (Bekker-Jensen *et al*, 2017; Wang *et al*, 2019). However, these studies did not provide quantitative information at the cell-type level. In comparison to bulk liver tissue, there are relatively few primary liver cells in isolates from an individual, requiring more efficient sample preparation procedures and more sensitive mass spectrometry acquisition methods. To address these challenges, we adopted state-of-the-art technologies - including the loss-less high-pH reversed-phase fractionation (Kulak *et al*., 2017). This allowed us to quantify the largest cell type-resolved quantitative human liver proteome atlas, consisting of 10,528 proteins assembled from 158,670 sequence-distinct peptides. The largest prior efforts investigated primary cells and only at one time point, identifying 6,788 proteins (Sun *et al*., 2010) and 9,791 proteins (Ölander *et al*, 2020).

Data independent acquisition (DIA) in MS-based proteomics has demonstrated superior performance in recent years in terms of proteome depth, data completeness and quantification reproducibility as compared to data dependent acquisition (DDA), and has become the preferred method in clinical studies (Bruderer *et al*., 2015; Guo *et al*, 2015; Hansen *et al*, 2021; Karayel *et al*, 2020). To achieve an optimal data quality, a deep and high-quality peptide spectral library is typically generated for a specific type of tissue. However, this step is laborious and not every group is equipped with the necessary instrumentation and technology to generate high-quality spectral libraries. The extensive dataset generated in our study can serve as a spectral library for high-throughput proteomic analysis of patient samples. With an optimized DIA method integrated with an advanced signal-processing algorithm termed “phased spectrum deconvolution” (ΦSDM) (Grinfeld *et al*., 2017), and Mq.Live for direct instrument control (Wichmann *et al*., 2019), we achieved proteome coverage of 6,000 proteins in 100 min measurement time in single-runs, demonstrating suitability of this workflow for analyzing clinical samples.

We investigated multiple dimensions of the human liver proteome: (i) four isolated primary liver cell types and three tissue types; (ii) dynamic proteome profiles during primary cell culture; and (iii) changes in the liver proteome of a cohort of patients with NASH and end-stage liver cirrhosis. The resulting quantitative data and its in-depth bioinformatics analysis highlighted new quantitative aspects of basic cell and tissue biology. For example, we estimate - using the “proteomic ruler” approach (Wisniewski *et al*., 2014) – that half of the protein mass in hepatocytes is composed of enzymes and about 1.2 of 4 billion protein molecules in each cell carry out metabolic pathway functions, reflecting their extremely high metabolic activity. Furthermore, there are 15-77 million protein copies in OXPHOS complexes I to IV in hHEPs, up to six-fold higher than other cell types. In contrast, less than 20% of protein mass in hLSECs is dedicated to enzymatic function, but significantly more to transporters, ribosomal proteins, cytoskeleton and extracellular matrix proteins, reflecting higher transcytosis and the functional demands for regulating sinusoidal blood flow (Poisson *et al*, 2017).

These cell-type level quantitative information can also help elucidate disease mechanisms and provide useful information for drug development. Our copy number catalogue contains quantitative information of disease-relevant proteins, such as those known to carry SNPs predisposing or protecting against NAFLD, including PNPLA3, GCKR and HSD17B13 (all most abundant in hHEPs). MBOAT7 is a transmembrane protein involved in phosphatidylinositol remodeling, and its rs641738C>T variant, strongly contributes to pathogenesis of a range of liver diseases in human genetic studies (Buch *et al*, 2015; Thabet *et al*, 2016; Thabet *et al*, 2017). We find its copy numbers to be 25 fold-higher in hHSCs compared to hHEPs. In view of the key role of hHSC in fibrogenesis, this raises interesting mechanistic questions. Among therapeutically relevant proteins, TGF-beta receptor and PDGF receptor have highest levels in hHSCs but are also present in all other liver cell types examined. Knowledge of cell type specific abundance should be useful for cell type-specific targeting, which could be more effective than global approaches with less adverse effects (Klein *et al*., 2019; Klein *et al*., 2012). Furthermore, our set of more than 100 proteins uniquely detected in one of the cell types, may serve as in developing targeted drug delivery methods or as new markers for isolating liver cell types. We found that cross-referencing the bulk and single-cell RNA sequencing data as well as the image collection generated by the Human Protein Atlas project helps to validate cell type specificity of the markers in a scalable manner.

Human primary cell culture are common systems for drug evaluation and development (Eglen & Reisine, 2011), but researchers need to know how close they mimic the in vivo situation, especially after culturing. In this regard, we previously observed significant changes in the proteome of murine primary hepatocytes upon cell culture (Azimifar *et al*., 2014). Consistent with this, we here observed extensive proteome drift upon cell culture with 41% hepatocyte proteins changing significantly in seven days. It is characterized by reduced levels of metabolic or energy homeostatic processes, including PPAR signaling – a pathway regulating lipid metabolism and inflammation, and an anti-NASH drug target under active development (Boeckmans *et al*., 2019; Gross *et al*, 2017; Wu *et al*., 2020). These changes should be considered when evaluating drug efficacy in cell models. Investigating dynamic proteome changes upon HSC activation can help in the development of therapies against liver fibrosis (Zhang *et al*, 2017). Our time-course experiment revealed the dynamics of this process, including the upregulation of proteins involved in ECM organization and immune response. Our finding of upregulated immune response may indicate a new role of HSC in promoting inflammation and immune responses in liver disease.

To connect our findings from human tissue and dynamic primary cell culture directly to patients, we investigated the liver proteome changes in NASH and cirrhosis in a cohort of 45 individuals. Previously, we have observed dramatic remodeling in the plasma proteome of patients with liver cirrhosis, whereas only a few proteins significantly changed in NAFLD (Niu *et al*., 2019). In line with this, only a handful proteins in the liver significantly changed in NASH, whereas half of the total liver proteome was significantly altered in liver cirrhosis, including increased levels of signaling pathways, extracellular matrix components and immunological response, as well as decreased levels in various metabolic pathways. This agrees with the fact that NASH is an intermediate but progressive form of chronic liver disease (Simon *et al*, 2021), whereas liver cirrhosis is the common end-stage of a wide variety of chronic liver diseases, characterized by substantial structural changes and impaired liver functions. This important insight agrees with the reversibility of NASH and underlines that therapeutic approaches for this stage may be more successful than in cirrhosis, in which the proteomic and thereby structural changes are very pronounced. Large portions of NAFLD and NASH do not present with liver-related events and this paper strengthens the rationale for approaches to improve the large regenerative capacity still available in NASH patients. By contrast cirrhosis has a very high liver-related mortality, rendering regeneration of the liver very improbable in the late stages, when complications of cirrhosis such as refractory ascites and ACLF occur. Our data and analyses are also valuable for cirrhosis research. For instance, the lack of important secretory capacity may point to therapeutic opportunities aiming at replacement of vital substances such as haptoglobin. This has only been done for albumin (Fernández *et al*, 2020) and coagulation factors (Bedreli *et al*, 2016) but not for haptoglobin. Our detailed catalogue of significantly altered proteins and pathways in the disease cohort and the primary cell time-course experiment provide a valuable tool for the research community to interpret data and discover new therapeutic targets. As an example, many of the proteins upregulated in bulk cirrhotic liver tissue can be mapped to hepatic stellate cell activation, highlighting intervention points before full blown cirrhosis. This can be used in conjunction with circulating biomarkers identified by proteomics such as the ones we previously showed for NAFLD and ALD in plasma (Niu *et al*., 2019; Niu *et al*., 2020).

To facilitate interactive data visualization of the large proteomics data sets, we built a dashboard application using the open source Dash framework in Python. We have also integrated pathological proteome changes in liver and plasma, which we generated in previous studies, allowing users to explore proteome changes at the level of both proteins and KEGG pathways. We hope that these data will become a valuable resource for basic, translational and clinical research focusing on liver pathophysiology, biomarker discovery and drug development.

## Materials and Methods

### Resources availability

#### Lead contact

Further information and requests for resources and reagents should be directed to and will be fulfilled by the Lead Contact, Matthias Mann.

#### Data and code availability

The mass spectrometry proteomics data have been deposited to the ProteomeXchange Consortium via the PRIDE (Vizcaíno *et al*, 2014) partner repository with the dataset identifier PXD027722. The dataset which is currently private can be accessed through the following reviewer account: Username. Password: SbZfXpMs. Summary statistics and bioinformatics results were provided in the supplementary tables. The code generated in this study have been uploaded to the Github repository https://github.com/llniu/Human_Liver_Proteome.

### Experimental model and subject details

#### Cell lines

Snap-Frozen cells were thawed in a water bath at 37°C and transferred to culture medium. Viability was controlled and was systematically over 95%. For cryopreservation, cells were centrifuged at 200 x g for 5 min. LX2 cells and TWNT-4 cells were grown with cell culture medium (DMEM + 20% FCS + Penicillin/Streptomycin) in 250 ml plastic flasks in a humidified 5% CO_2_ incubator at 37°C. After reaching 80% confluency, cells were passaged with a 1:3 split ratio. Detachment was achieved by incubating the cells with 0.05% trypsin/EDTA solution (solved in PBS) for 5 min at 37°C. HepG2 cells were grown with cell culture medium (DMEM + 10% FCS + Penicillin/Streptomycin) in 250 ml plastic flasks in a humidified 5% CO_2_ incubator at 37°C. Cells were seeded at density of about 3 × 10^6^ cells/80 cm^2^. After reaching 80% confluency, cells were passaged with a 1:2 split ratio. SK Hep1 cells were cultured in a humidified 5% CO^2^ incubator at 37°C. Cells were grown in culture medium (M199 + 10% FCS + Penicillin/Streptomycin). Cells were plated at density of about 4 × 10^6^ cells/80cm^2^ in flasks coated with collagen IV. After reaching 80% confluency, cells were passaged with a 1:10 split ratio. Detachment was achieved by incubating the cells with 0.05% Trypsin/EDTA solution (solved in PBS) for 5 min at 37°C.

#### Primary cells

Primary human HSC, KC, HEP and LSEC were obtained from Samsara sciences (San Diego, CA, USA). The cells used in the comparative analysis of basal proteome between the four cell types have not been previously cultured after isolation. Characteristics and purity of the isolated cells were assessed by the provider (liver cell characterization attached as supplemental materials). For the time-course primary cell culture experiment, snap-frozen primary cells were thawed in a water bath at 37°C and transferred to culture medium. Viability was controlled and was systematically over 95%. For cryopreservation, cells were centrifuged at 200 x g for 5 min. Primary human HSC were grown in cell culture medium (DMEM + 10% FCS + Penicillin/Streptomycin) in a humidified 5% CO^2^ incubator at 37°C. Cells were seeded at density of about 6,370 cells/cm^2^. After reaching 85% confluency, cells were passaged with a 1:3 split ratio. Detachment was achieved by incubating the cells with 0.05% Trypsin/EDTA solution (solved in PBS) for 5 min at 37°C. Primary human KC were grown in cell culture medium (RPMI 1640 + 10% FCS + Penicillin/Streptomycin) in a humidified 5% CO^2^ incubator at 37°C. Cells were plated at a concentration of 0.3 × 10^6^ cells/ml. Primary human HEP were grown in cell culture medium (Williams’ Medium E + 1% ITS + 10-7M Dexamethasone + Penicillin/Streptomycin + 10 mM HEPES) in a humidified 5% CO^2^ incubator at 37°C. Primary human LSEC were grown in cell culture medium (EBM-2 + EGM-2 singlequots) in a humidified 5% CO^2^ incubator at 37°C. Cells were plated at a seeding density of about 3,000 cells/cm^2^. Cells were plated in flasks coated with collagen I.

#### Human tissues

Liver tissue, hepatic artery and portal vein which were used to generate the fractionated, deep proteomes were collected from three receivers and three donors of liver transplantation during 2001 and 2003. For the human cohort (N=45 in total), additional 10 liver samples from patients with cirrhosis requiring liver transplantation (performed between 2001 and 2003) were used. Patients with liver cirrhosis were classified as Child-Pugh class B (n=4) or C (n=6) with a median score of 10 and had a median MELD (model of end-stage liver disease) score of 18.5 (minimum 13, maximum 23). Liver samples of patients with morbid obesity and non-alcoholic steatohepatitis (NASH, n=20), as well as samples from obese but liver-healthy individuals (n=10) were collected during bariatric surgery, which was performed at the Department of Bariatric, Metabolic and Plastic Surgery, St. Franziskus-Hospital Cologne, Germany, between July 2018 and May 2019 (Rheinwalt *et al*, 2020). Additional five liver samples from healthy donors of liver transplantation (performed between 2001 and 2003) were used as healthy controls, adding to 15 in total. Patients with NASH had a median fibrosis stage of F2 (minimum F1, maximum F3) (Kleiner *et al*, 2005), a median NAFLD activity score (NAS) of 6 (minimum 5, maximum 7 on a scale of 0 to 8) and had at least perisinusoidal fibrosis. Liver fibrosis stage from biopsy: F1/2/3 = 4/14/2. NAFLD activity score: 5/6/7 = 3/15/2. Liver steatosis score: 1/2/3 = 1/14/5. Hepatocyte ballooning score: 1/2 = 1/19. Lobular inflammation score: 1/2 = 4/16. Obese but liver-healthy control individuals had less than 5% of parenchymal steatosis. The diagnosis was performed independently by two experienced pathologists as described elsewhere (Schierwagen *et al*., 2020). Samples were washed with ice-cold PBS, snap-frozen in liquid nitrogen and stored in -80°C after collection. The investigation was approved by the ethical committee of the University of Bonn (document no. 029/13 and 194/17, respectively) by the ethics committees of the regional Medical Association Nordrhein (project identification code 2017110) in accordance with the declaration of Helsinki. All patients signed an informed consent before being enrolled in the study.

### Method details

#### Sample preparation for MS analysis

Tissue samples were ground to a frozen powder using a mortar and pestle in liquid nitrogen. Powdered samples were then resuspended in 350 μl of sodium deoxycholate (SDC) reduction and alkylation buffer (PreOmics GmbH, Martinsried, Germany) and boiled for 10 min while vortexing at 1200 rpm to denature proteins (Kulak *et al*, 2014). The lysates were sonicated at full power for 30 cycles with 30s intervals using a water bath sonicator (Diagenode Bioruptor®, Liège, Belgium). Protein content was determined by Tryptophan assay. An aliquot of 150 μl homogenate was digested overnight with LysC and trypsin in a 1:50 ratio (μg of enzyme to μg of protein) at 37°C and 1700 rpm. On the following day, boiling and sonicating was repeated followed by an additional step of digestion for 2h (1:100 ratio). Peptides were acidified to a final concentration of 0.1% trifluoroacetic acid (TFA) to quench the digestion reaction. Peptide concentration was estimated using Nanodrop and 20 μg of peptides was loaded on two 14-gauge Stage-Tip plugs. Peptides were washed first with isopropanol/1% TFA (200 μl) and then 0.2% TFA (200 μl) using an in-house-made Stage-Tip centrifuge at 2000 x g. Peptides were eluted with 60 μl of elution buffer (80% acetonitrile/1% ammonia) and dried at 60°C using a SpeedVac centrifuge (Eppendorf, Concentrator plus). Dried peptides were redissolved and sonicated in 5% acetonitrile/0.1% TFA, and concentration was measured using Nanodrop. About 50 μg of peptides were fractionated into 8 fractions using basic reverse phase high-pH fractionation with the SPIDER fractionator (PreOmics GmbH, Martinsried, Germany). Cell lines, isolated and cultured human primary cells were processed similarly to the tissue samples without liquid nitrogen crushing.

#### LC-MS/MS

All samples were measured using LC-MS instrumentation consisting of an EASY-nLC 1200 system (Thermo Fisher Scientific, San Jose, CA) interfaced on-line with a Q Exactive HF-X Orbitrap or Orbitrap Exploris 480 equipped with a FAIMS Pro Interface (Thermo Fisher Scientific, Bremen, Germany). The latter LC-MS instrumentation setup was used in the time-course experiment of primary cell culture. For all samples, purified peptides were separated on 42.5 cm HPLC-columns (ID: 75 µm; in-house packed into the tip with ReproSil-Pur C18-AQ 1.9 µm resin (Dr. Maisch GmbH)). For each LC-MS/MS analysis, around 0.5 µg peptides were injected for the 100 min gradients. Peptides were loaded in buffer A (0.1% formic acid) and eluted with a linear 82 min gradient of 3-23% of buffer B (0.1% formic acid, 80% (v/v) acetonitrile), followed by a 8 min increase to 40% of buffer B. The gradients then increased to 98% of buffer B within 6 min, which was kept for 4 min. Flow rates were kept at 350 nl/min. Re-equilibration was done for 4 μl of 0.1% buffer A at a pressure of 980 bar. Column temperature was kept at 60 °C using an integrated column oven (PRSO-V2, Sonation, Biberach, Germany).

MS spectra of fractionated samples were acquired with a Top15 data-dependent MS/MS scan method (DDA, topN method). Target values for the full scan MS spectra was 3e6 in the 300-1,650 m/z range with a maximum injection time (IT) of 25 ms and a resolution of 60,000 at m/z 200. Precursor ions targeted for fragmentation were isolated with an isolation width of 1.4 m/z, followed by higher-energy collisional dissociation (HCD) with a normalized collision energy of 27 eV. Precursor dynamic exclusion was activated with 30s duration before triggering the subsequent scan. MS/MS scans were performed at a resolution of 15,000 at m/z 200 with an automatic gain control (AGC) target value of 1e5 and an IT of 25 ms.

MS spectra of unfractionated patient samples were acquired in single-shot with a data-independent acquisition (DIA) method, enabled by MaxQuant.Live (Wichmann *et al*., 2019) in which the scan protocol was defined. Each acquisition cycle was consisted of a survey scan at resolution of 60,000 with an AGC of 3e6 and IT of 100ms, followed by 66 DIA cycles at resolution of 15,000 with an AGC of 3e6 and IT of 22ms at range 300-1650 m/z (Table S7). HCD fragmentation was set to normalized collision energy of 27%. In all scans, PhiSDM (Grinfeld *et al*., 2017) was enabled with 100 iterations, spectra type was set to centroid.

For spectra acquisition in the time-course experiment of primary cell culture, a FAIMS Pro Interface was mounted between the electrospray ionization source and the mass spectrometer (Orbitrap Exploris 480). The ion source was set to a voltage of 2650 (V) in positive ion mode with an ion transfer tube temperature of 275°C. FAIMS mode was set to ‘standard resolution’ with a total carrier gas flow of 4.6 L/min throughout the entire acquisition period. For single-shot analysis, MS spectra were acquired using DIA mode with intra-analysis compensation voltage (CV)-switching. Each acquisition cycle was consisted of a survey scan at resolution of 120,000 with a normalized AGC target (%) of 300% and 28 ms of injection time at scan range of 350-1650 m/z, followed by 22 DIA cycles at resolution of 15,000 with a normalized AGC target (%) of 3000% and 25ms of injection time repeated for three CVs (−40V, -55V, and -70V), totaling a cycle time of around 3s (Table S8). HCD fragmentation was set to normalized collision energy of 30%. To create a spectra library based on gas-phase fractionation, a pooled, unfractionated primary cell sample for each cell type was analyzed in single-shots using seven methods of different CVs stepping from -40V to -70V with an increment of -5V. These gas-phase fractionation methods consisted of a survey scan at resolution of 120,000 followed by 66 DIA scans at resolution of 15,000 (Table S8). The resulting MS spectra were analyzed together with data acquired by the above-described single-shot method of intra-analysis CV switching to boost identifications.

#### MS data processing

All raw files of fractionated samples were analyzed by MaxQuant v.1.5.3.30 software (Cox & Mann, 2008) using the integrated Andromeda Search engine (Cox *et al*, 2011) and searched against the Uniprot human database (April 2017 release including isoforms and sequence variants). Enzyme specificity was set to trypsin with a maximum of two missed cleavages. The search included cysteine carbamidomethylation as fixed modification and oxidation on methionine and N-terminal acetylation as variable modifications with a minimum length of seven amino acids. A false discovery rate (FDR) of 1% was set to PSM and protein levels. The ‘match between runs’ algorithm was activated to transfer MS/MS identifications between runs where applicable (Nagaraj *et al*, 2012). Label-free quantification was performed with the integrated MaxLFQ algorithm using a minimum ratio count of 2 (Cox *et al*., 2014). A spectral library was generated from the fractionated samples for single-injection DIA analysis of patient samples in the clinical cohort (N=45).

All raw files of patient samples were analyzed by Spectronaut (version 13.3) with default settings except that the normalization strategy for ‘cross-run normalization’ was wet to ‘local normalization’ based on rows with “Qvalue complete” (Bruderer *et al*., 2015). A FDR of 1% was set to peptide precursor level and 1% to protein level. The FDR method of Storey was used (Storey & Tibshirani, 2003). The library generated from fractionated samples described above was used in the targeted analysis of single-shot DIA data against the human Uniprot fasta database (January 2018 release including isoforms and sequence variants).

All raw files in the time-course experiment of primary cell culture were analyzed by Spectronaut (version 14) in directDIA mode against the human Uniprot fasta database (January 2018 release including isoforms and sequence variants). Default settings were used except that the normalization strategy for ‘cross-run normalization’ was wet to ‘local normalization’ based on rows with “Qvalue complete”.

For the proteome comparison across four primary cell types (fractionated samples), we filtered proteins quantified in this study for at least two valid values in three biological replicates of at least one cell type, resulting in 8,866 proteins. For the patient samples, we filtered proteins quantified in the cohort for at least 70% valid values at experimental group level (healthy, NASH and cirrhosis group), resulting in 5,843 proteins with an average of 5,382 proteins per sample and an overall data completeness of 91.8%. For comparison between time points of the same cell type in the time-course experiment of primary cell culture, we filtered proteins for at least two valid values in three biological replicates of at least one time point within each cell type, resulting in total number of proteins ranging between 6,100 and 6,900 proteins in the three cell types. The missing values were replaced by drawing random samples from a normal distribution (down-shifted mean by 1.8 standard deviation (SD) and scaled SD (0.3) relative to that of proteome abundance distribution, with which we performed the statistical analysis.

#### Bioinformatics and statistical analysis

Statistical and bioinformatics analysis was performed with the Perseus software (version 1.6.2.1) and Python software. One-way ANOVA significance level was controlled with FDR below 5% with Benjamini-Hochberg for multiple hypothesis testing. GO Enrichment Analysis in the primary cell type proteome was performed with ClueGo (Bindea *et al*, 2009), a plug-in app in Cytoscape (Shannon *et al*, 2003), with default settings except the following changes: Ontologies file of Biological Process was downloaded from EBI-Uniprot on September 04, 2018 which contains 15,947 terms with 17,940 available unique genes. Customed reference set which contains 9,223 unique genes (quantified in this study) was used in Fisher’s exact test. Term significance was corrected by Benjamini-Hochberg with a FDR of below 1%. GO tree levels was controlled at 2-3 for HEP and KC, and 2-4 for LSEC and HSC with a threshold of 10 genes and 10% of genes per term to maximize the information presentable. Both GO term fusion and grouping are activated. GO Enrichment Analysis in patient samples between NASH, cirrhosis and control was performed with the online Gene Ontology Resource (geneontology.org). All python scripts used to generate corresponding results and figures are available at the GitHub repository https://github.com/llniu/Human_Liver_Proteome. The interactive dash board application was built using the open source web application framework of Plotly Dash and Python. Source code can be found in the same GitHub repository.

## Acknowledgments

We thank members of our groups in Copenhagen and Munich for help and discussions. We also thank Jeppe Madsen, Martin Rykær, Parvaneh Sadeghi, Johannes Mueller, Simone Schopper, Gudrun Hack and Silke Bellinghausen for excellent technical assistance. We also thank the funding agencies: the Challenge Programme *MicrobLiver* funded by Novo Nordisk Foundation (grant No. NNF15OC0016692), the Max Planck Society for the Advancement of Science, the European Union’s Horizon 2020 research and innovation program (grant agreement on 686547 to the MSmed project; 825694 to the MicrobPredict project), the Novo Nordisk Foundation for the Clinical Proteomics group (grant NNF15CC0001), and the Novo Nordisk Foundation for the Copenhagen Bioscience PhD Program (NNF16CC0020906).

## Author Contributions

L.N. acquired and interpreted the proteomics data, designed and developed the project and wrote the manuscript. P.L.G. designed the project, performed proteomic sample preparation and edited the manuscript. R.G. prepared proteomics sample and edited the manuscript. A.S. guided the data analysis and revised the manuscript. F.M. optimized proteomic acquisition method. S.D. performed proteomic sample preparation. N.J.W.A critically reviewed and edited the manuscript. S.K., C.O., F.E.U. and R.S. performed cell culture and edited the manuscript. J.T. designed the project, provided all biological and clinical materials, and edited the manuscript. M.M designed and supervised the project, interpreted the data and edited the manuscript.

## Figure legends

**Fig EV7** | **Liver proteome remodelling under pathological conditions. c**. Volcano plot of the *P*-values vs. the log2 protein abundance differences in cirrhosis compared with control. Significance level is controlled by *P*-value (independent two-sample *t*-test, two-sided) and minimum fold change (s0 parameter in Perseus) indicated by the cutoff curve, highlighting proteins related to ECM organization (red-colored), majority of which up-regulated and proteins associated with fatty acid beta-oxidation (blue-colored), majority of which down-regulated. **d**. Volcano plot of the P-values vs. the log2 protein abundance differences in NASH compared with control, highlighting significantly up- and down-regulated proteins (color-coded in red and blue respectively).

## Expanded View Figure legends

**Fig EV1.**
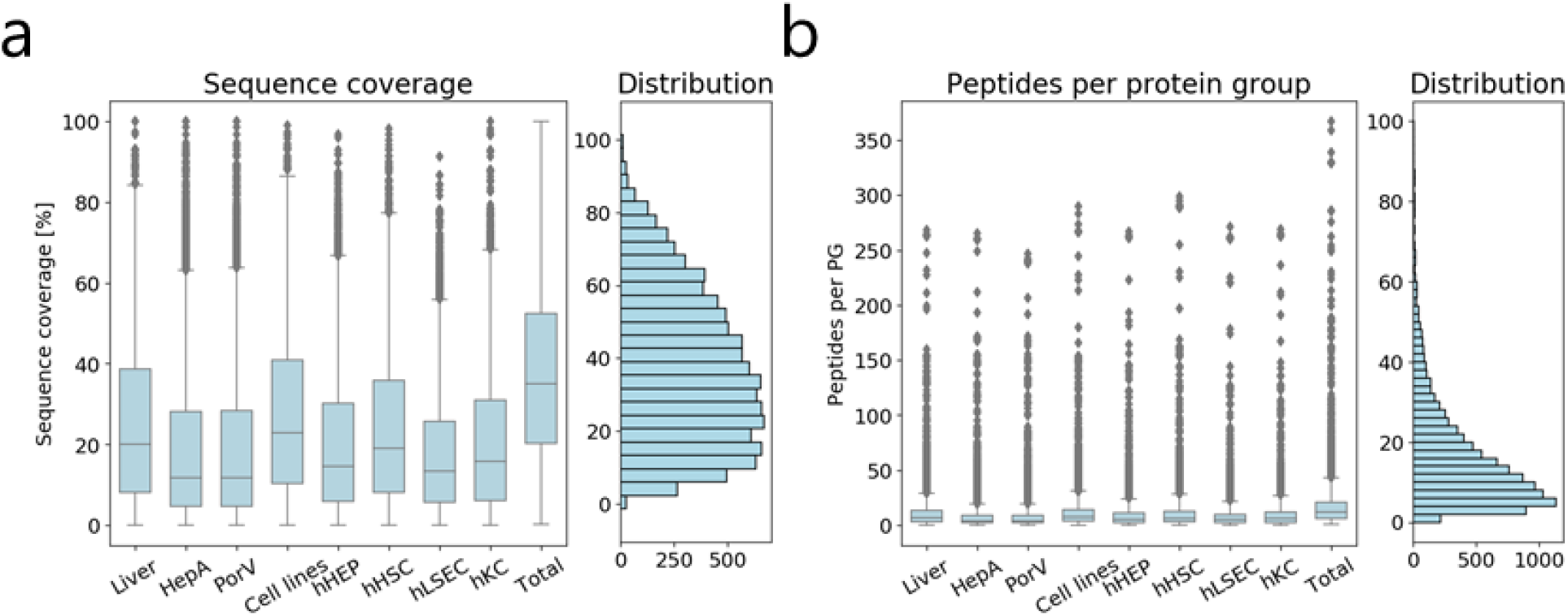
Proteomics data overview. **a**. Box plots of sequence coverage distribution of all proteins quantified in each tissue/cell type, and in all samples combined. **b**. Box plots of number of identified peptides per protein group in each tissue/cell type, and in all samples combined.

**Fig EV2.**
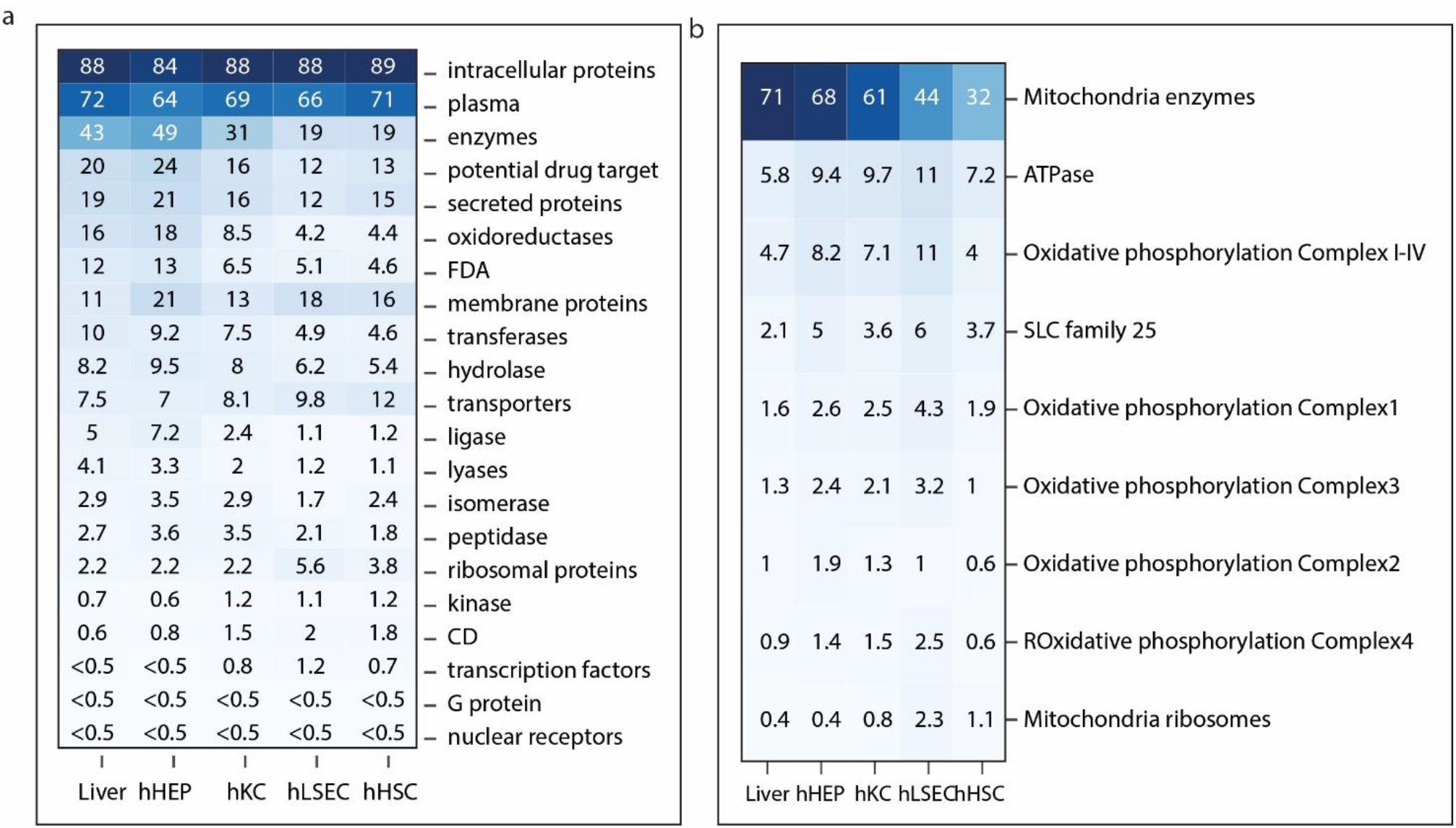
Cellular mass composition based on protein classes. **a**. Percent protein mass by protein classes. Protein class annotation was based on the HPA classification. **b**. Percent protein mass in the mitochondria by selected protein classes. Oxidative phosphorylation is the sum of Complex 1-4.

**Fig EV3.**
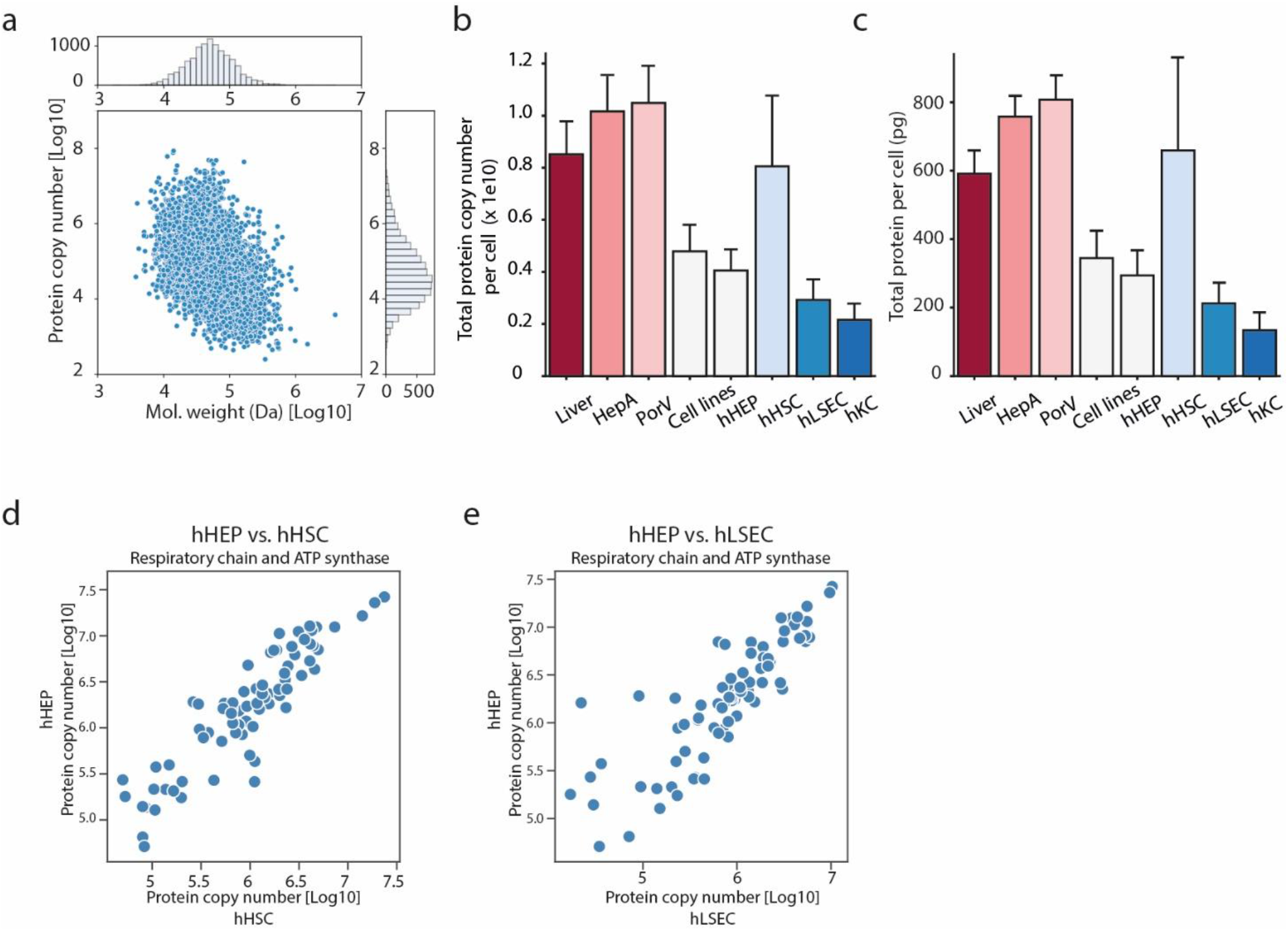
Protein copy number estimation. **a**. Protein copy number versus molecular weight in the hepatocyte proteome, with histogram on the x- and y-axis representing distribution of the molecular weights and protein copy numbers, respectively. **b-c**. Total protein copy number (b) and total protein mass per cell (c) in liver tissues and cell types estimated by the ‘proteome ruler’ approach. **d-e**. Protein copy number for members of the oxidative phosphorylation complex I-V in hepatocytes and hepatic stellate cells (d), sinusoidal endothelial cells (e).

**Fig EV4.**
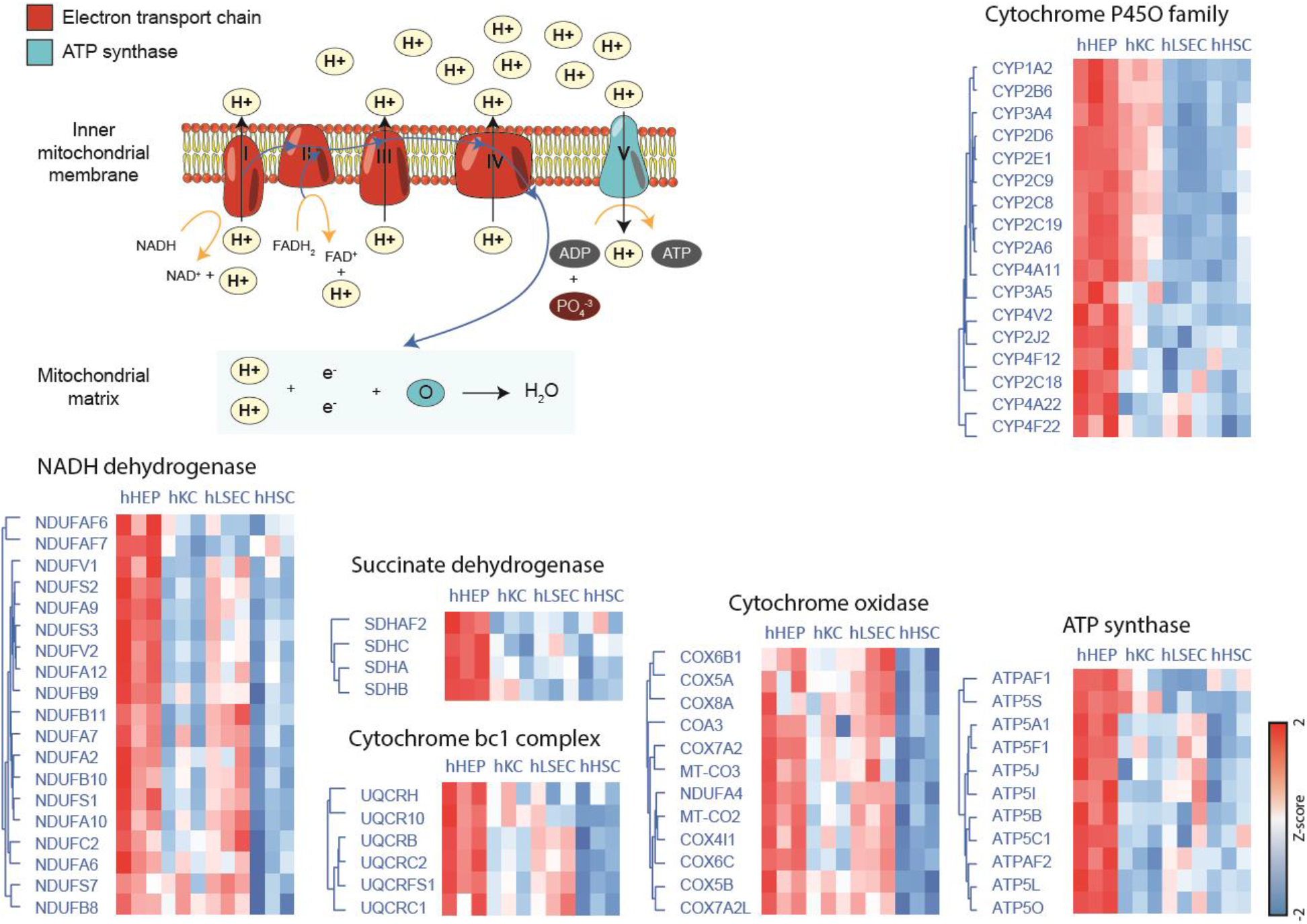
Hepatocyte enriched proteins in respiratory chain and drug metabolism. Proteins comprised the electron transport chain and ATP synthase were extracted and clustered based on their Z-score normalized abundance across cell types.

**Fig EV5.**
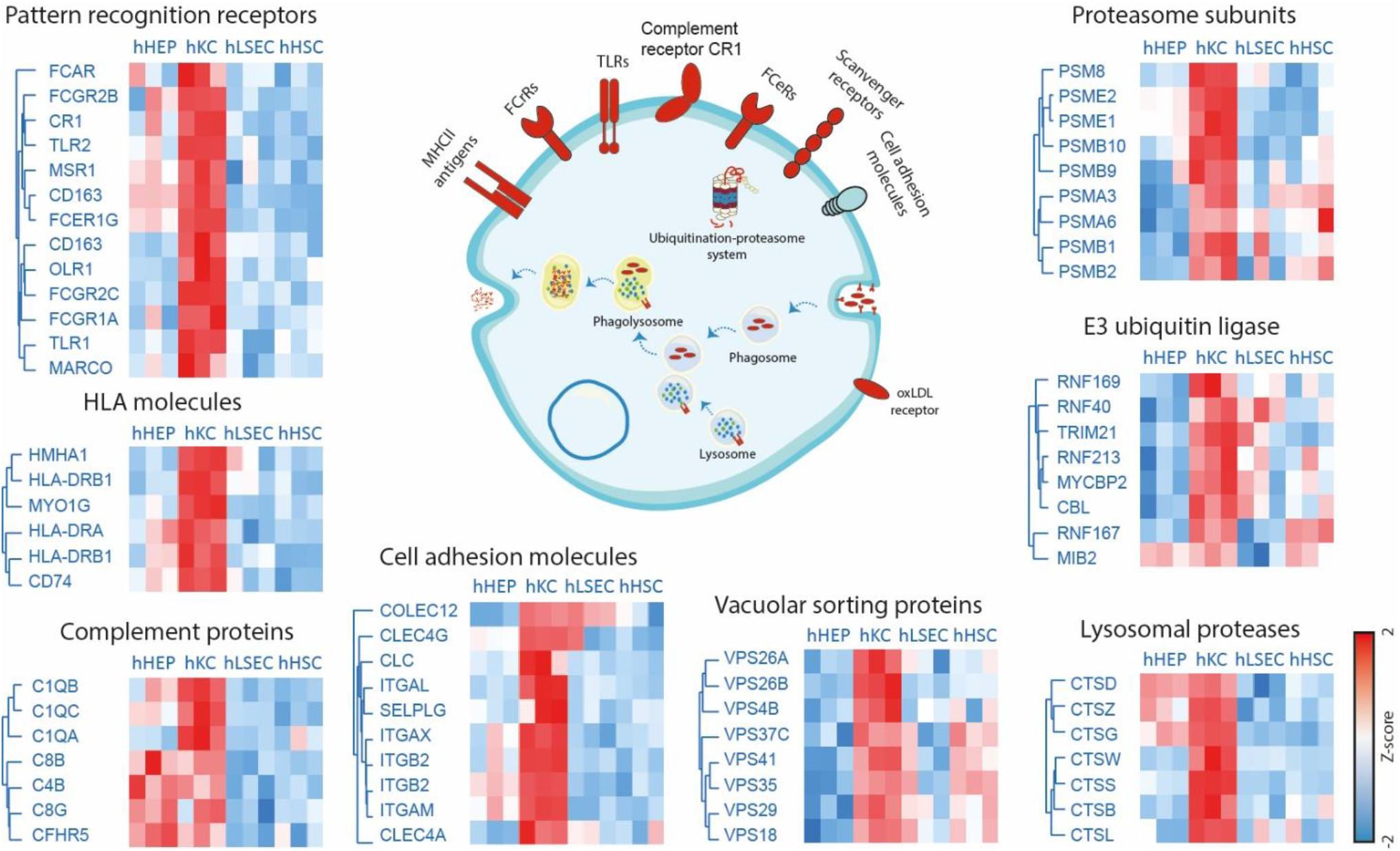
Kupffer cell enriched proteins in antigen processing and presentation pathway. Proteins related to the immune response and protein degradation were extracted and clustered based on their Z-score normalized abundance across cell types.

**Fig EV6.**
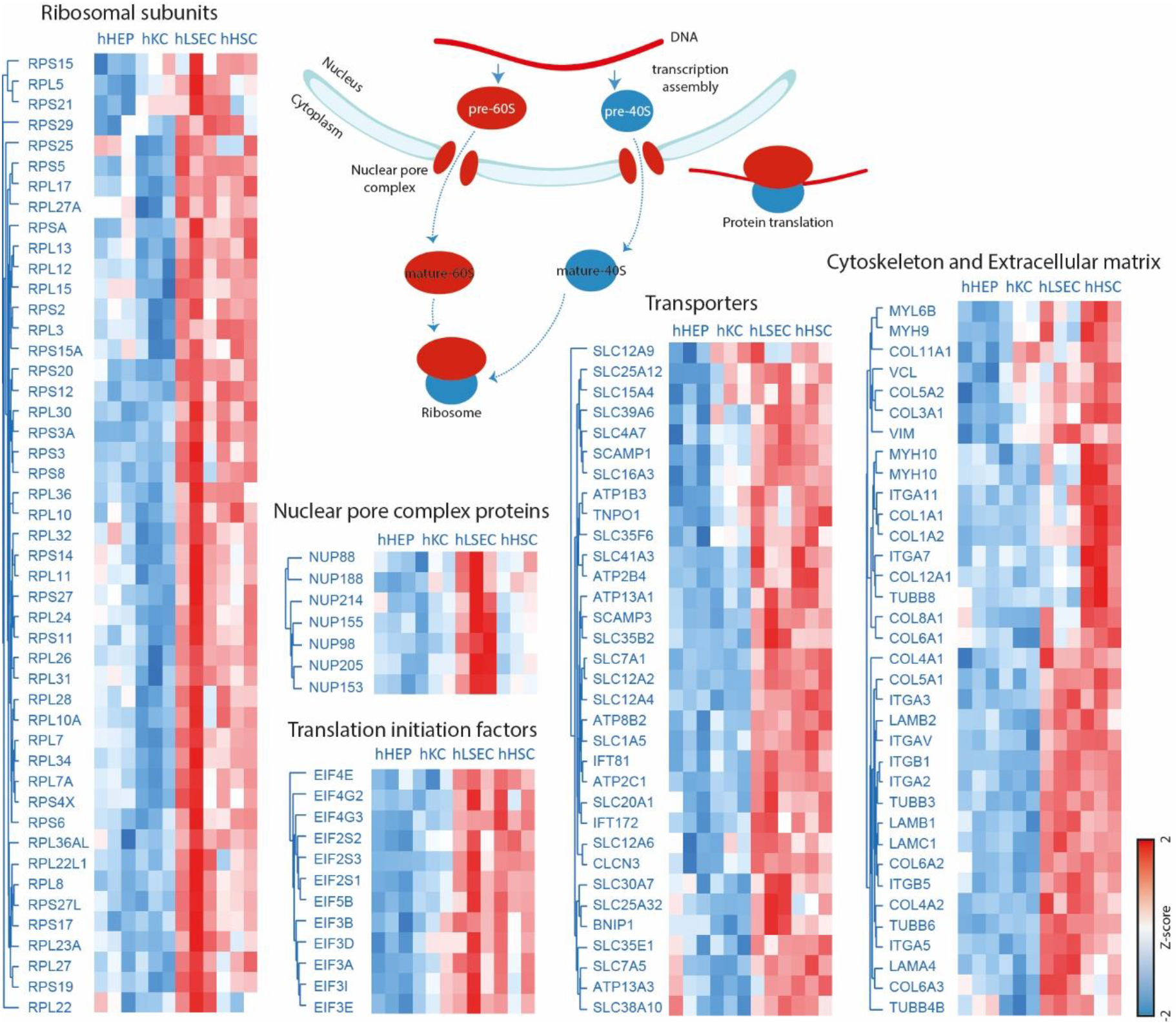
LSEC-enriched proteins in ribosomal biogenesis and translation & HSC-enriched proteins in cytoskeleton and extracellular matrix. Proteins were clusterd based on their Z-score normalized abundance across cell types.

